# Probing variability in a cognitive map using manifold inference from neural dynamics

**DOI:** 10.1101/418939

**Authors:** Ryan J. Low, Sam Lewallen, Dmitriy Aronov, Rhino Nevers, David W. Tank

## Abstract

Hippocampal neurons fire selectively in local behavioral contexts such as the position in an environment or phase of a task,^1-3^ and are thought to form a cognitive map of task-relevant variables.^1,4,5^ However, their activity varies over repeated behavioral conditions,^6^ such as different runs through the same position or repeated trials. Although widely observed across the brain,^7-10^ such variability is not well understood, and could reflect noise or structure, such as the encoding of additional cognitive information.^6,11-13^ Here, we introduce a conceptual model to explain variability in terms of underlying, population-level structure in single-trial neural activity. To test this model, we developed a novel unsupervised learning algorithm incorporating temporal dynamics, in order to characterize population activity as a trajectory on a nonlinear manifold—a space of possible network states. The manifold’s structure captures correlations between neurons and temporal relationships between states, constraints arising from underlying network architecture and inputs. Using measurements of activity over time but no information about exogenous behavioral variables, we recovered hippocampal activity manifolds during spatial and non-spatial cognitive tasks in rats. Manifolds were low-dimensional and smoothly encoded task-related variables, but contained an extra dimension reflecting information beyond the measured behavioral variables. Consistent with our model, neurons fired as a function of overall network state, and fluctuations in their activity across trials corresponded to variation in the underlying trajectory on the manifold. In particular, the extra dimension allowed the system to take different trajectories despite repeated behavioral conditions. Furthermore, the trajectory could temporarily decouple from current behavioral conditions and traverse neighboring manifold points corresponding to past, future, or nearby behavioral states. Our results suggest that trial-to-trial variability in the hippocampus is structured, and may reflect the operation of internal cognitive processes. The manifold structure of population activity is well-suited for organizing information to support memory,^1,5,14^ planning,^12,15,16^ and reinforcement learning.^17,18^ In general, our approach could find broader use in probing the organization and computational role of circuit dynamics in other brain regions.

---

The activity state of a neural circuit can be described as a point in a high-dimensional coordinate system, termed the state space, where each coordinate represents the firing rate of an individual neuron. Neural population activity over time can be viewed as a trajectory through this state space (Fig. 1a, “population activity”). Underlying properties of the network and its inputs can confine this trajectory to a low-dimensional, possibly nonlinear subspace, termed a manifold, with a much fewer number of intrinsic coordinates that encode cognitive and behavioral variables. For example, hippocampal attractor models predict that position in the environment is encoded in this manner during spatial navigation^19^ and mental exploration,^15^ and similar models of computation and memory have been proposed for many systems.^20-25^ As a subspace of state space, the manifold delimits the allowable joint activity patterns, manifesting as correlations between neurons. For example, in Fig. 1a, neuron 1’s instantaneous firing rate can be perfectly predicted from neurons 2 and 3. Furthermore, the manifold can constrain the temporal evolution of neural activity, as continuous transitions between distant states must move through intermediate states along the manifold. Therefore, restriction to a low-dimensional manifold implies that individual neurons are not free to vary independently; rather, their activity reflects underlying, population-level structure. In this case, trial-to-trial variability would correspond to variation in the underlying network trajectory, which must remain on the manifold. Experimentally testing this picture requires a means to reconstruct a manifold from population recordings, which should ideally 1) discover nonlinear structure, 2) capture temporal relationships between states, 3) recover topological and geometric properties of the manifold, and 4) provide explicit mappings between neural activity and coordinates on the manifold. Powerful methods have been developed for uncovering low-dimensional, latent structure in data,^26-38^ but do not generally meet all criteria simultaneously.

**Figure 1:**
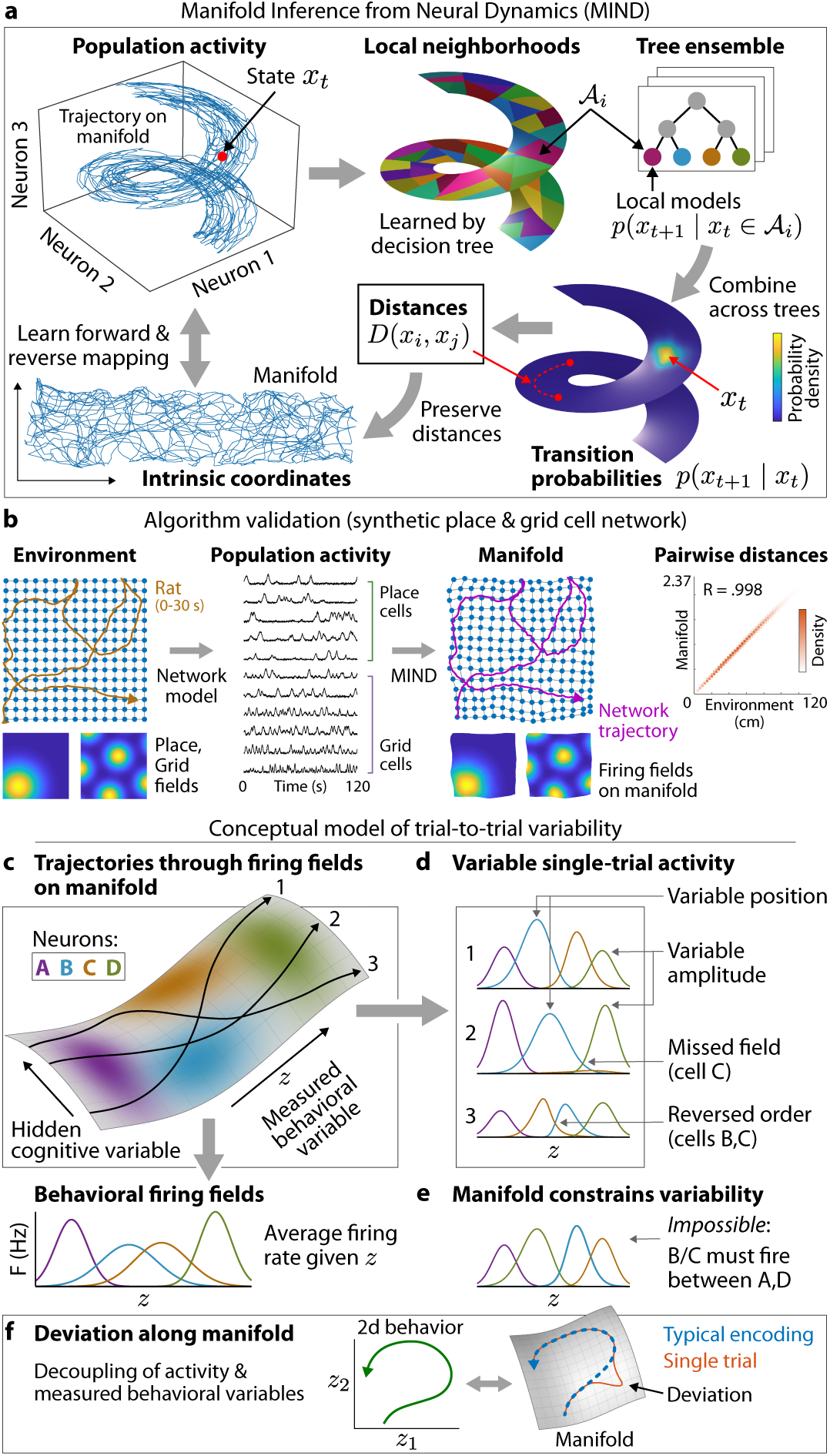
Manifold discovery and model of trial-to-trial variability. **a,** Schematic of manifold discovery algorithm (see Methods for details). Shown for a synthetic example with three neurons. Activity forms a random walk in firing rate space, constrained to a 2d spiral manifold. **b,** A 2d, nonlinear manifold is recovered from the activity of 100 place and grid cells as a simulated rat explores a 2d arena (subset of cells and time points shown). The network’s trajectory on the manifold matches the rat’s trajectory in the environment. Points plotted on the manifold correspond to population activity at the grid of locations shown in the environment (plots scaled and rotated to align for visual comparison). Lines connect points that are neighbors in the environment; the manifold preserves spatial topology. Pairwise distances on the manifold and environment are correlated (R=.998, 3.1 × 10^6^ pairs of points). Cells form firing fields on the manifold that match their firing fields in the environment. **c-f,** Conceptual model of trial-to-trial variability. **c,** Single-trial population activity corresponds to trajectories on a nonlinear manifold (shown for 3 trials). Directions along the manifold represent internal variables that may correspond to behavioral measurements or be hidden (one of each type shown). Neurons fire as a function of network state, forming firing fields on the manifold (firing rate shown as color intensity for four cells). Behavioral firing fields average out the hidden variable. **d,** The extra dimension (hidden variable) allows the network to take different paths through manifold firing fields despite identical behavioral conditions. Consequently, cells fire with different amplitude, position, and order across trials. **e,** The manifold constrains trial-to-trial variability. For example, local topology determines the permitted sequences in which cells can fire. **f,** Trial-to-trial variability can also arise if activity decouples from instantaneous behavioral conditions and the network moves independently along the manifold. Shown for a 2d behavioral trajectory.

To reconstruct a manifold while satisfying the above criteria, we developed a new approach: Manifold Inference from Neural Dynamics (MIND; Fig 1a, see Methods). We view the manifold as an abstract object representing constraints on population activity over time. It is embedded in a high-dimensional state space but may be intrinsically low-dimensional, in which case the population activity can be described by a small number of intrinsic coordinates. In applying MIND, we first learned transition probabilities between states, using a novel algorithm based on a randomized ensemble of decision trees. Each tree adaptively partitioned state space into local neighborhoods, with a continuous probability distribution over future states for each neighborhood. After combining estimates across trees, we used the transition probabilities to construct a distance metric between states, which defined their proximity on the manifold. Distances had the property that they were smaller for pairs of states that transitioned to each other with higher probability, through a smaller number of intermediate states. We obtained intrinsic manifold coordinates representing the network trajectory by embedding states into a low-dimensional Euclidean space such that distances were approximately preserved. We learned functions mapping neural activity space to intrinsic manifold coordinates (and vice versa), which are useful in evaluating how well the manifold captures the neural activity, and in explaining the variability of this activity across trials. MIND distances can also be studied directly, making it possible to estimate geometric and topological properties of the manifold, and enabling the reconstruction of more general manifolds such as circles and spheres.

We validated our method on synthetic data from a simple model of the hippocampal-entorhinal system during spatial navigation. The model contained 50 place cells and 50 grid cells^1^ that fired as a simulated rat explored a 2d arena. Each cell fired according to its place field or grid field, which specified firing rate as a function of the rat’s position (Fig. 1b). Because the rat moved uniformly throughout the arena and each position strictly determined a unique firing pattern, activity should lie on a manifold that is isometric to the physical environment up to an overall scale. Accordingly, we fit a 2d manifold to the population activity using MIND. As expected, the recovered manifold was geometrically equivalent to the environment, and the network’s trajectory on the manifold matched the rat’s trajectory in the environment (Fig 1b). Computing firing rates as a function of manifold coordinates yielded firing fields on the manifold. In contrast to place and grid fields, which are a direct function of behavior, manifold firing fields relate individual neuronal activity to the underlying network state. Because of the correspondence between network states and the environment, each neuron’s firing field on the manifold matched its place or grid field in the environment (Fig. 1b).

Unlike the synthetic data, real population activity may not be entirely determined by measured behavioral variables. Neural circuit function is often characterized by estimating behavioral firing fields (e.g. place and grid fields), which represent the average activity of single neurons over repeated behavioral conditions. However, despite the often strong relationship between average firing and behavior, neural activity has been widely observed to vary across repeated trials.^6-10^ This variability could correspond to noise, or might represent additional cognitive information.^6,11-13^ In the latter case, we hypothesize that additional cognitive variables manifest as correlated structure in the population activity, in the same manner as measured behavioral variables. In particular, they could be encoded by additional dimensions of a manifold.

In this conceptual model (Fig. 1c), population activity lies on a nonlinear manifold, as described above. Movement along the manifold reflects the internal dynamics of the system, which may be modulated by sensory, behavioral, or cognitive processes. Each intrinsic dimension of the manifold represents an internal variable, which may correlate with a behavioral parameter that the experimenter has measured, or may be hidden (e.g. an unmeasured parameter, or cognitive variable with no external correlate). Individual neurons fire as a function of the overall network state— forming firing fields on the manifold—and the system takes varying trajectories through these fields over time. In contrast, behavioral firing fields represent expected firing rates given the measured variables, and are defined by averaging out all other sources of variability. We consider two complementary ways in which structured trial-to-trial variability can arise. First, the extra, hidden dimensions allow the system to take different trajectories through fields on the manifold even if behavioral conditions are fixed. Individual neurons may thus fire with different amplitudes and in different behavioral conditions on each trial (Fig. 1d). They may even fire in different order, or fail to fire when the the trajectory completely misses a field on the manifold. However, the manifold constrains the variability that can occur. For example, the manifold’s local topology limits the possible sequences in which cells can fire (Fig. 1e). Second, trial-to-trial variability can also arise if population activity temporarily decouples from current behavioral conditions, as might occur during spontaneous memory, prediction, or mental exploration events.^5,12,14-16,39^ In our model, such deviations from the typical encoding correspond to the network moving along the manifold independently of measured behavioral variables (Fig. 1f).

To test our conceptual model, we analyzed population activity recorded from the dorsal CA1 hippocampal region of 4 rats. In addition to a standard spatial task (random foraging task, RFT), the rats performed a novel sound manipulation task^3^ (SMT) where they remained in a fixed position, and pressed a lever to adjust an auditory tone to a goal frequency (Fig. 2a). Individual neurons had localized behavioral firing fields on the environment (RFT) or sound frequency (SMT), which collectively represented the measured behavioral variables.^3^ For our analyses, we created population activity vectors from the firing rates of simultaneously recorded cells, which we analyzed separately for each behavioral session. We studied 6 sessions in total, 3 from each task, with 15-52 neurons per session.

**Figure 2:**
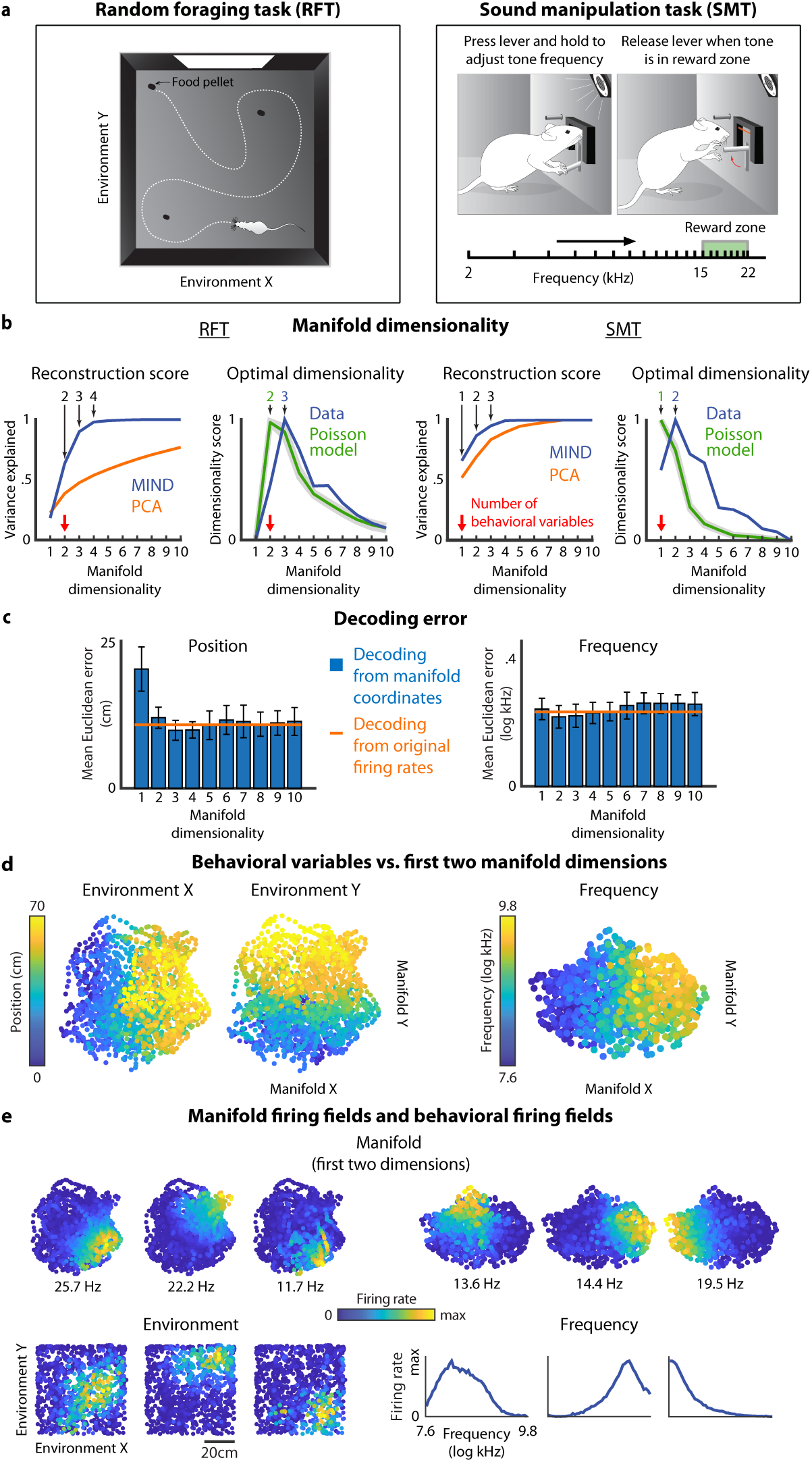
Properties of hippocampal activity manifolds. Analyses shown for rat 103 during one RFT session (*left column*) and one SMT session (*right column*). **a**, Depiction of task structure. **b**, *Left:* Fraction of variance in the population activity explained by manifolds of increasing dimensionality, constructed by MIND. In comparison, the fraction of variance explained by optimal linear manifolds as constructed by PCA. *Right:* Intrinsic dimensionality score vs. manifold dimensionality. We used the peak of the intrinsic dimensionality score to select an optimal dimensionality. In comparison to a Poisson null model whose activity was determined by the measured behavioral variables, the hippocampal activity had an extra dimension. Gray band indicates standard error across synthetic Poisson datasets. **c**, Decoding error when predicting behavioral variables from intrinsic manifold coordinates of increasing dimensionality, or from activity vectors containing the original firing rates. Error bars represent s.d. over cross-validation folds. **d**, Representation of behavior along the first two coordinates (ordered by variance) of the optimal manifold (RFT 3d, SMT 2d, as estimated in **b**). Each datapoint is colored by the values of the behavioral variables at the corresponding timepoint. **e**, *Top:* Manifold firing fields for 3 cells from each task, depicting firing rate as a function of manifold coordinates. Visualized along the first two coordinates of the optimal manifold, as in **d**. Each cell’s maximum firing rate is indicated. *Bottom:* Behavioral firing fields for the same cells, depicting firing rate as a function of measured behavioral variables. For visual comparison, the manifold plots in **d,e** are aligned to the behavioral coordinates by rotation and scaling.

To assess whether the hippocampal activity was constrained to a low-dimensional manifold, we used MIND to fit manifolds of increasing dimensionality. For each choice of dimensionality *d*, we mapped activity vectors to d-dimensional intrinsic manifold coordinates, which we then mapped back to activity space to define reconstructed activity vectors; these represent the projection of the original activity onto the manifold fit by MIND. We computed a reconstruction score, defined as the fraction of variance in the activity vectors explained by the reconstructed activity vectors, which was closer to 1 when the activity was closer to the manifold. The reconstruction score saturated at a low dimensionality across behavioral sessions in both tasks (.91 ± .05 by d = 3; mean ± s.d.; Fig. 2b, Extended Data Figs. 1 and 2), indicating that the hippocampal population activity lies near a low-dimensional manifold. To control for overfitting, we computed cell prediction scores^27^ (where each manifold was used to predict the activity of held-out cells), which saturated or attained their optimum at low dimensionalities (Extended Data Fig. 3). To test whether the reconstructed manifolds were nonlinear, we compared MIND to principal component analysis (PCA), which recovers linear manifolds with optimal reconstruction score (Fig. 2b, Extended Data Figs. 1 and 2). MIND attained greater reconstruction scores at lower dimensionality, particularly for the RFT, where PCA required an order of magnitude more dimensions to reach the same performance. As a further control to ensure that low-dimensional manifold fits were not inevitable, we verified that high-dimensional simulations with properties matched to the data were not well-fit at low dimensionalities (Extended Data Fig. 4). Thus, in both tasks, our results suggest that hippocampal population activity is restricted to lie near a low-dimensional, nonlinear manifold.

To estimate the intrinsic dimensionality of the hippocampal activity, we fit manifolds at each dimensionality d, as above. For each d, we computed an intrinsic dimensionality score (IDS, see Methods). We took the peak of the IDS curve to define the optimal dimensionality, and we validated this estimator on simulated datasets (Extended Data Fig. 4). For the hippocampal activity, the optimal manifold dimensionality in each task exceeded the number of measured behavioral variables (RFT, d = 3 for all rats, SMT, d = 2 for rats 103 and 106, d = 3 for rat 107; Fig. 2b, Extended Data Figs. 1 and 2). This suggests that the behavioral variables do not completely determine the neural activity, which varies along a greater number of dimensions. As a control, we analyzed synthetic population activity from a Poisson null model,^6^ where each neuron’s activity was generated according to its behavioral firing field, and did not contain any sources of variability beyond Poisson noise. In this case, we found neural activity manifolds whose dimensionalities matched the measured behavioral variables (d = 2, RFT, d =1, SMT; Fig. 2b). Therefore, in contrast to the variability in the data, Poisson variability is not sufficient to increase the dimensionality of the manifold, providing further evidence that the neural variability represents structure rather than noise. In summary, our analyses indicate that the hippocampal activity is well-fit by nonlinear manifolds of dimension much less than the number of neurons. However, for each task, the activity manifold has at least one extra dimension, suggesting that the neural activity is not completely determined by the measured behavioral variables, and may encode additional information.

To characterize the relationship between the manifold and behavior, we first attempted to decode the rat’s instantaneous position in the RFT or sound frequency in the SMT from the intrinsic manifold coordinates at each timepoint (Fig. 2c, Extended Data Figs. 1b, 2b). Decoding performance at the optimal manifold dimensionality (identified as above) was comparable to that achieved with the original firing rates (mean Euclidean decoding error with manifold coordinates vs. firing rates: 11 vs. 11 cm for RFT, .28 vs. .32 log kHz for SMT; mean across rats). Thus, the manifold captures all information about measured behavioral variables that is available to the decoder in the raw firing rates. Decoding performance was similar for manifolds fit at a lower dimensionality, equal to the the number of behavioral variables *d*_*b*_ (Fig. 2c, Extended Data Figs. 1b, 2b). Because manifold dimensions are non-redundant, this suggests that the extra dimension we identified above carries little information about measured behavioral variables. Rather, it represents additional information in structured population activity.

Position and sound frequency were encoded smoothly along the manifold, and change in each variable corresponded to continuous movement along a different direction on the manifold (Fig. 2d, Extended Data Figs. 1c, 2c). To quantify this relationship, we compared distances between corresponding pairs of points when calculated using the *d*_*b*_-dimensional behavioral coordinates versus the first *d*_*b*_ manifold coordinates with greatest variance. Pairwise distances in both spaces were correlated, indicating geometric similarity between the manifold and behavior (Pearson correlation .70 ± .06 for RFT and .55 ± .16 for SMT; mean ± s.d. across rats). Thus, the manifold captures a map-like representation of behavioral variables in the neural activity, and MIND can recover this representation in an unsupervised manner. In comparison, related manifold learning algorithms (isomap and PCA) did not reproduce this geometric structure, and preserved less information about behavior (Extended Data Fig. 5, Methods section 18).

Many neurons formed compact firing fields on the manifold, indicating that they fired preferentially in a local neighborhood of network states, as in our conceptual model (Fig. 1c, 2e, Extended Data Figs. 6, 7). Field positions remained stable when manifolds were fit separately to each half of the behavioral session (Extended Data Fig. 8), suggesting that manifold firing fields represent genuine constraints on the relationship between cell activity and network state that persist throughout the local behavioral context. When viewed along the first *d*_*b*_ manifold coordinates, these fields largely recapitulated the cells’ localized behavioral firing fields, as expected given the geometric similarity of the behavioral space and the manifold along these dimensions (Fig. 2e, Extended Data Figs. 6, 7). However, as above, the manifold contained an extra dimension beyond the measured behavioral variables. Many cells’ firing fields were localized and stable along this extra dimension as well, suggesting that their firing reflects additional structure beyond the encoding of measured behavioral variables (Fig. 2e, Extended Data Figs. 6, 7).

Could this extra dimension account for trial-to-trial variability, as in our conceptual model? In the SMT, the rat swept through the same sound frequencies on every trial, and many cells had frequency-tuned behavioral firing fields.^3^ However, they fired at a different sound frequency and with a different number of spikes on each trial (Fig. 3a). We quantified this variability by examining single-trial activity as a function of sound frequency, and estimating the normalized position and amplitude of the activity peak on each trial. The interdecile range (IDR) across trials measured the position and amplitude variability of each cell, ranging from 0 (no variability) to 1 (maximum variability) (Fig. 3b). As previously observed in spatial tasks,^6^ many cells exhibited high variability in the SMT (position IDR .56 ± .25, amplitude IDR .69 ± .11, mean ± s.d. for all cells across rats). Position variability was broadly distributed across cells, whereas nearly all cells had highly variable peak amplitudes.

**Figure 3:**
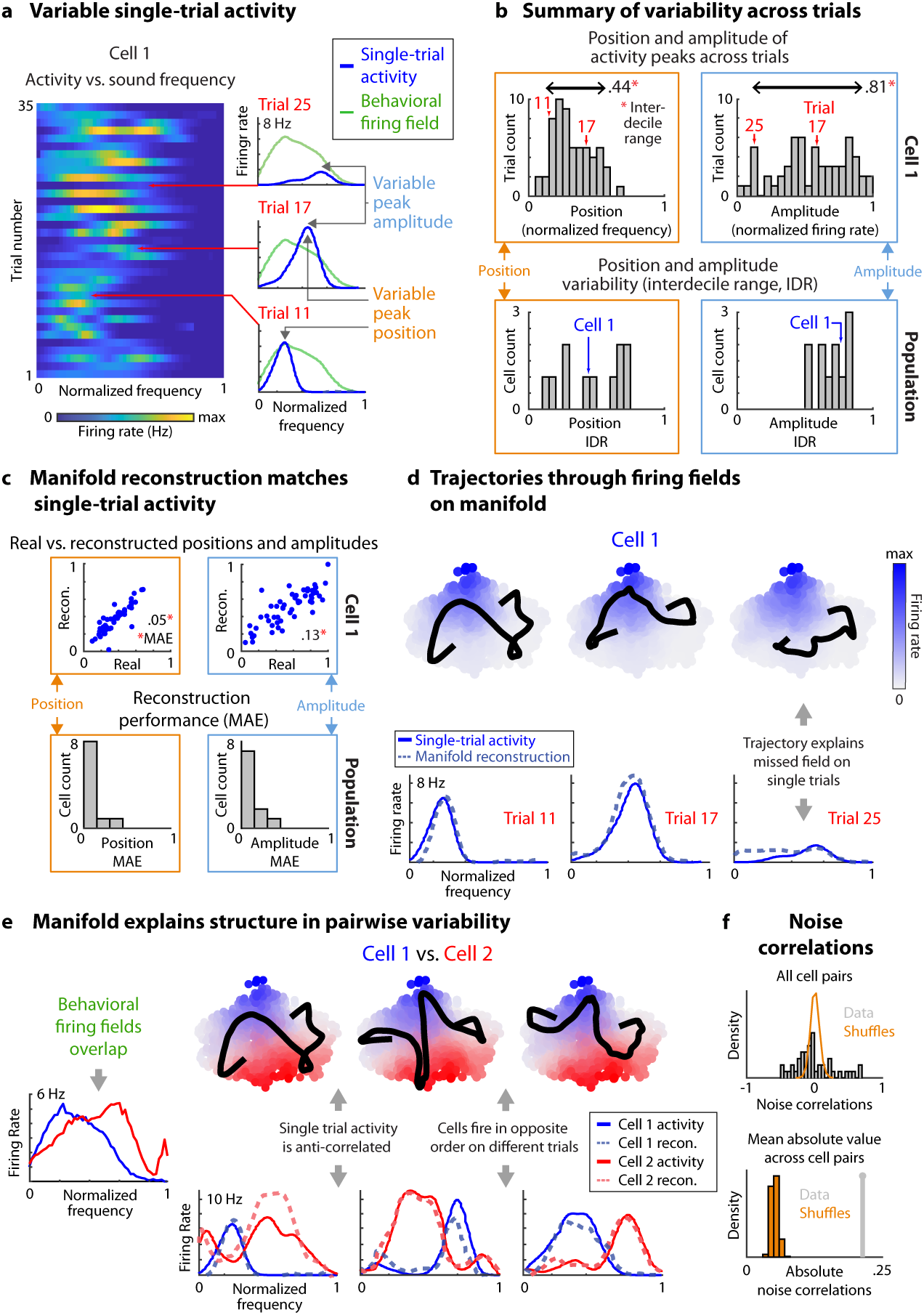
Manifold trajectories explain trial-to-trial variability in the sound manipulation task. Analyses shown for rat 103. **a,** *Left:* Single-trial activity as a function of sound frequency for an example cell (cell 1). Frequency is normalized to range from 0 to 1 on a log scale. First 35 (out of 59) trials are shown. *Right:* Activity vs. frequency for 3 example trials. The activity peak occurred at a different position and amplitude on each trial. **b**, *Top:* Distribution of positions and amplitudes of activity peaks across trials, for cell 1. Amplitudes are scaled to the cell’s maximum firing rate. The interdecile range (IDR) of each distribution summarizes trial-to-trial variability. An IDR closer to 1 indicates greater variability. *Bottom:* Position and amplitude IDRs for all analyzed cells (those with sufficiently high firing rate). **c**, Reconstructed activity (i.e. the projection of real activity onto the 2d manifold) captured trial-to-trial variability. *Top:* Amplitude and position for real vs. reconstructed activity peaks on each trial, for cell 1. As a summary of reconstruction performance, the average absolute difference between real and reconstructed amplitudes and positions across trials (mean absolute error, MAE) is indicated. *Bottom:* MAE between real and reconstructed positions and amplitudes, for all analyzed cells. **d**, *Top:* Network trajectory on the 2d manifold for 3 example trials. Overlaid on cell 1’s manifold firing field, depicted here as reconstructed activity vs. manifold coordinates. The network’s path through the firing field determined the reconstructed activity for cell 1. *Bottom:* Real and reconstructed activity vs. sound frequency on each example trial. **e**, *Left:* Behavioral firing fields for cells 1 and 2. *Right:* Network trajectory on the 2d manifold for 3 different example trials, overlaid on the manifold firing fields for each cell. In contrast to behavioral firing fields, the manifold fields were nearly disjoint. Correspondingly, the single-trial activity of the two cells was anti-correlated, and their order of firing could vary. **f**, Variability on the manifold (as in e) can lead to noise correlations between cells, defined as pairwise correlations not predicted from the frequency tuning of each cell. *Top:* Distribution of noise correlations across cell pairs. Shuffled datasets preserve behavior-related correlations but destroy noise correlations (see Methods). *Bottom:* Mean absolute value of noise correlations across cell pairs, compared to the same shuffles as above.

Despite this variability, the recovered manifold successfully predicted single-trial activity. As above, activity reconstructions accounted for most of the variance in the raw neural activity (Fig. 2b). Furthermore, in the SMT, single-trial peak positions and amplitudes in the reconstructed activity matched those in the original data (Fig. 3c) (mean absolute error .09 ± .06 for position, .14 ± .08 for amplitude; mean ± s.d. across cells and rats). Thus, trial-to-trial variability is not arbitrary. Rather, it corresponds to variation in the trajectory of the entire population along a low-dimensional manifold. As in our conceptual model (Fig. 1c,d), the single-trial activity of individual neurons reflected the interaction of this trajectory with firing fields on the manifold. In particular, the existence of an extra dimension beyond sound frequency allowed activity to vary over repetitions of the same frequency (Fig. 3d,e). For example, cells could fail to fire at their preferred frequencies when the 2d trajectory traversed around, rather than through their firing fields on the manifold-something impossible for a one-dimensional system (Fig. 3d). This phenomenon also occurred for the 3d RFT manifold (Extended Data Fig. 9). Furthermore, despite the monotonic increase in frequency on every trial, cells with adjacent firing fields on the manifold could fire in different order, depending on which field the trajectory entered first (Fig. 3e).

Finally, the fact that population activity is intrinsically low-dimensional but higher-dimensional than the behavior suggests that trial-to-trial variability is correlated across cells. Such correlated variability is commonly referred to as noise correlation, although it may reflect underlying structure, such as fluctuation in internal states.^7^ When population activity is viewed as a trajectory on a manifold capturing these internal states, noise correlation arises from the arrangement of firing fields on the manifold. For example, the cells shown in Fig. 3e fire at similar sound frequencies on average because their manifold firing fields are aligned along the first dimension, which encodes frequency. However, these fields are largely non-overlapping, and are arranged along the second dimension such that a trajectory avoiding one field must necessarily pass through the other. Therefore, given a fixed sound frequency, trial-to-trial variation of the trajectory along the second dimension will produce anti-correlated fluctuations in the two cells’ activity. To quantify this effect, we estimated noise correlations between pairs of cells in the SMT using standard procedures (see Methods). We tested the null hypothesis of zero noise correlation using a permutation test where permutations destroyed noise correlations, but preserved behavior-related correlations arising from the frequency tuning of each neuron. All rats had cell pairs with negative and positive noise correlations exceeding those in the permuted data (Fig. 3f, Extended Data Fig. 2d). The mean absolute noise correlation across cells was significantly greater than zero (.14 ± .07, mean ± s.d. across rats, *p* < .004 for each rat). Thus, trial-to-trial variability is correlated across cells, as expected given the manifold structure of population activity.

Our conceptual model describes a second means by which trial-to-trial variability could arise: temporary decoupling of neural activity from the current behavioral conditions (Fig. 1f). As above, there is a general correspondence between intrinsic manifold coordinates and measured behavioral variables (Fig. 2c,d). However, this correspondence could momentarily break if the network were to move independently along the manifold. To test this picture, we decoded the rat’s position in the RFT from the network’s 3d coordinates on the manifold. The predicted and actual trajectories through the environment were typically synchronized. However, the predicted trajectory could deviate during local episodes, in which network activity encoded future, past, or nearby locations in the environment (Fig. 4a). We refer to these deviations as leads, lags, and detours, respectively. Predicted and actual trajectories were more closely synchronized for a manifold fit to our Poisson null model (Fig. 4a), where activity encodes the rat’s current position by construction (as above).

**Figure 4:**
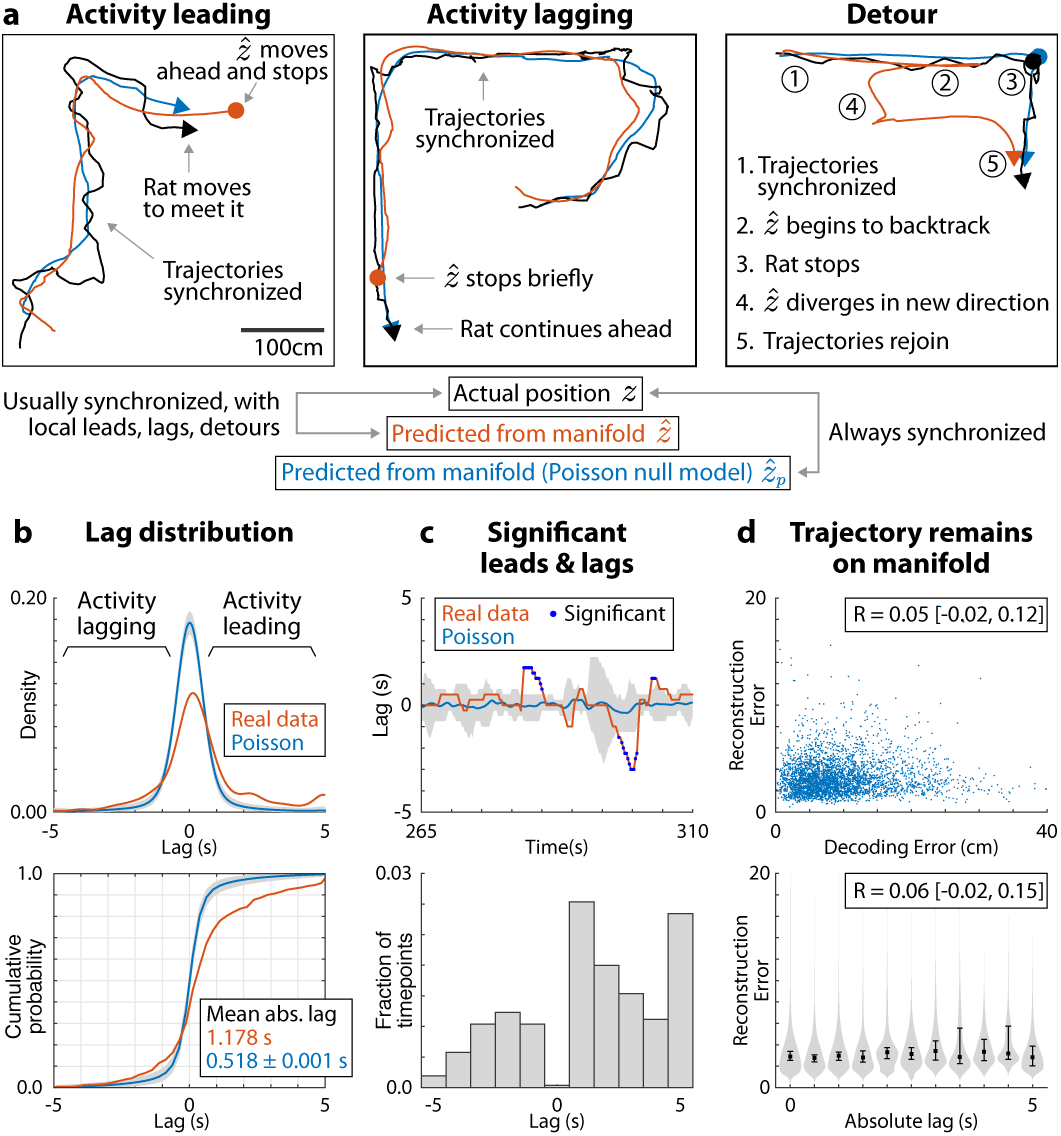
Local deviations from encoding current position. Analyses shown for rat 103. **a,** Example trajectories showing the rat’s position during random foraging and its predicted position given the network’s location on the manifold. Population activity typically encoded the rat’s current position, but could locally deviate to encode future (lead), past (lag), or nearby (detour) positions. Activity under the Poisson null model encoded instantaneous position by construction, producing tight coupling between actual and predicted position. **b,** Estimated time lag between the predicted and actual position. *Top*: Probability density function of lag aggregated across time (kernel density estimate). *Bottom*: Corresponding cumulative distribution function (empirical CDF). The gray region is a pointwise 95% confidence band and the light blue curve is the average across synthetic Poisson datasets. **c,** The hypothesis that lag in the real data exceeded that under the Poisson model was tested at each timepoint, controlling the false discovery rate at a level of .05. *Top*: Estimated lag at each timepoint, for an example interval. For the Poisson model, the gray region contains the 2.5 to 97.5 percentiles of the lag distribution at each timepoint, and the blue curve is the mean. *Bottom*: The relative frequency of statistically significant timepoints with each lag, expressed as a fraction of the total number of timepoints included in the analysis. **d,** The network did not depart from the manifold during leading, lagging, or detour deviations, as measured by the manifold reconstruction error. *Top*: Manifold reconstruction error vs. decoding error. *Bottom*: Manifold reconstruction error vs. absolute lag. Points and error bars show the median and 95% bootstrap confidence intervals. Gray regions (violin plot) show the conditional density of reconstruction error given absolute lag. Pearson correlation is shown with 95% bootstrap confidence intervals.

To quantify leading and lagging deviations, we used dynamic time warping to estimate the temporal lag between the predicted and true position at each timepoint. The distribution of lags over time was peaked around zero, indicating that the trajectories were typically synchronized. Tails of the distribution correspond to occasional leading and lagging deviations (Fig. 4b, Extended Data Fig. 10). The lag distribution for the Poisson model indicates the magnitude of leads and lags expected by chance. Lead and lag magnitudes, measured by the mean absolute lag, were significantly greater for the real data than the Poisson model (p < 5 × 10^−4^ for all rats, estimated by Monte Carlo simulation from the Poisson model). To identify statistically significant lead and lag events while controlling for local trajectory properties, we tested the hypothesis at each timepoint that absolute lag in the real data exceeded that in the Poisson model (false discovery rate controlled at the level .05). Significant lead and lag events were present in all rats, with leads occurring more frequently than lags (Fig. 4c, Extended Data Fig. 10). Rats also had significant detour events (see Methods), where the actual position differed strongly from the predicted position, and this difference could not be accounted for by lead or lag. Significant leads, lags and detours during random foraging behavior are illustrated in Supplementary Video 1 (see also Extended Data Fig. 11).

We next asked whether the network trajectory remains on the manifold during deviations from encoding the current position. If true, there should be no increase in the manifold reconstruction error, which measures the distance between an activity vector and its projection on the manifold (as described above). Deviations—where population activity encodes a location other than the rat’s current position—are characterized by an increase in position decoding error. However, reconstruction error and decoding error were essentially uncorrelated (Fig. 4d top, Extended Data Fig. 10c) (Pearson correlation 0.05, 95% CI [−0.02, 0.12] for the rat shown in Fig. 4; see Extended Data Fig. 10c for other rats). Similarly, there was no apparent relationship between reconstruction error and the magnitude of lag (Fig. 4d bottom, Extended Data Fig. 10d) (Pearson correlation 0.06, 95% CI [−0.02, 0.15] for the rat shown in in Fig. 4; see Extended Data Fig. 10d for other rats). Therefore, hippocampal activity does not depart from the manifold during leading, lagging, and detour deviations. Rather, the relationship between activity and behavior changes momentarily: during deviations, the network traverses states on the manifold that corresponded to future, past, or nearby behavioral states. An intriguing possibility is that leads, lags, and detours might reflect prediction, memory, or mental exploration processes in the hippocampus.^5,12,14-16^’^39^

Prior work^3,40^ suggests that task performance, both in spatial navigation and cognitive processes in general, activates a sequence of neural activity in the hippocampal formation in which firing fields are elicited parametrically with progress through behavior. Neighboring and partially overlapping fields represent the order and adjacency of behavioral states, useful in linking events in episodic memory,^1,5,14^ planning future actions,^12,15,16^ and reinforcement learning.^17,18^ Here we expand that conceptual picture by demonstrating that, in two different cognitive tasks, the population level activity in the hippocampus lies near relatively low-dimensional manifolds in the state space of neural activity. Previously observed firing field sequences correspond to movements through adjacent states on the manifold. Importantly, the manifold structure not only determines the classical tuning properties (behavioral firing fields) but the neural variability as well. In this new conceptual view, during random foraging in space,“place” cells do not fire because the animal enters the place field; they fire when the neural circuit traverses the manifold field. Single trial activity averaged across manifold traversals produce the behavioral firing fields, but different trajectories produce variability, reflecting internal dynamics of the system. Importantly, this variability is not random, but structured according to the manifold, producing noise correlations. In this view, neural coding and structured variability are cast in a common framework: both correspond to movement along the same manifold.

Large scale recording of neural activity at cellular resolution is being increasingly used to measure neural circuit dynamics at the population level during behavior. MIND adds to the emerging set of methods for characterizing this network activity as low-dimensional trajectories in state space and analyzing its properties.^26-38^ Our focus here has been on using MIND-based manifold characterization and analysis to probe the origin of variability in the hippocampus, but it should also be useful in characterizing neural dynamics in sensory perception,^8,23^ motor control^25,28^ and cognitive tasks such as working memory^20,21^ and integration of evidence during decision-making.^41^ Future goals include determining how general manifolds are in brain dynamics, how their geometric structure and topology depends on the areas recorded, how their properties evolve over time,^42^ and how specific forms of computation such as decision-making, memory recall, or action planning can be better understood in the conceptual framework of movement along or between manifolds—a general approach that may be thought of as “computing with manifolds.”

## Acknowledgements

We thank A. Akrami, R. Chaudhuri, C. Domnisoru, I. Fiete, J. Pillow, M. Schottdorf, and B. Scott for insightful discussions and comments on the manuscript. The illustrations in Fig. 2a are by J. Kuhl. This work was supported by the Simons Collaboration on the Global Brain.

---

**Extended Data Figure 1:**
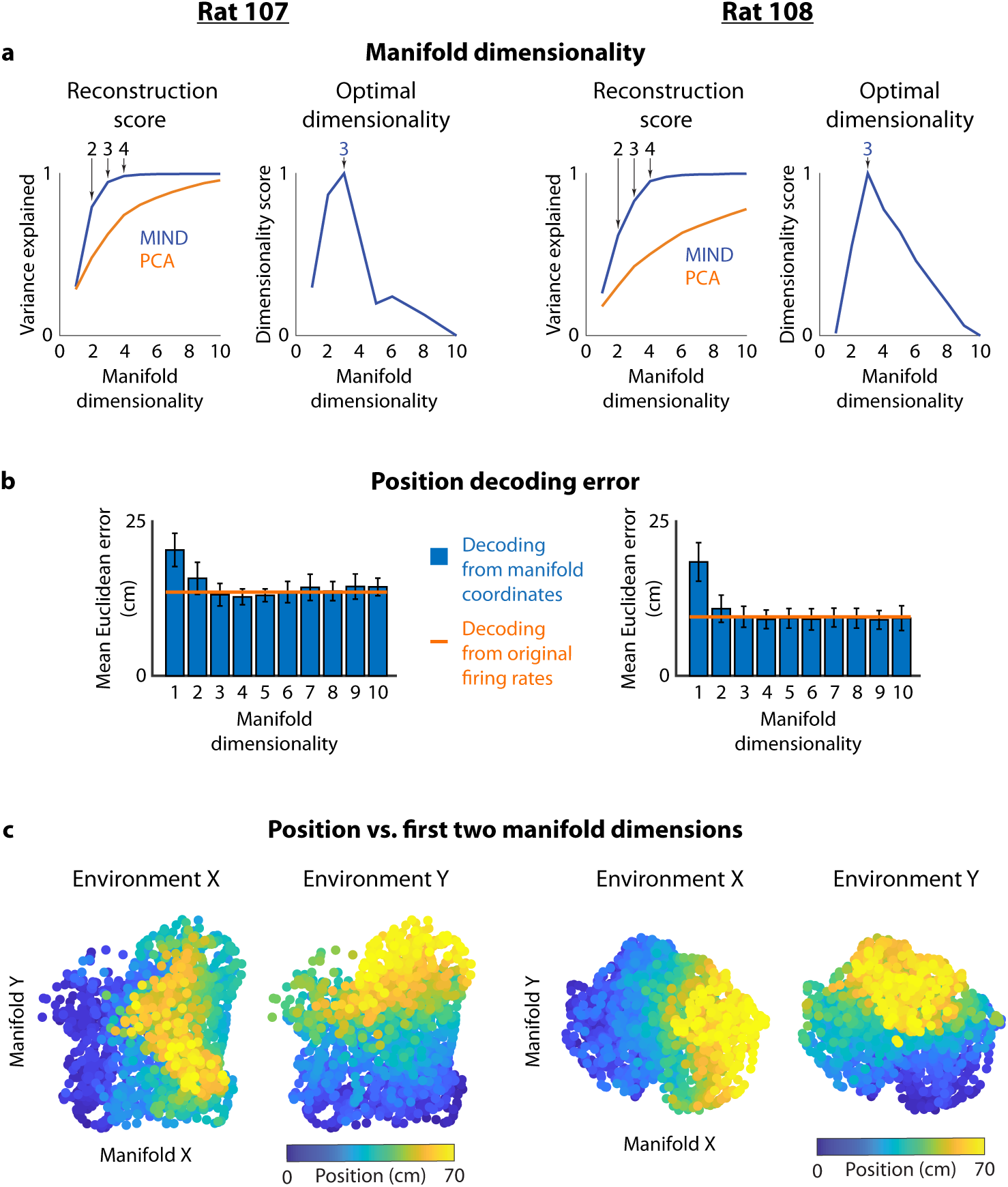
Properties of RFT manifolds for behavioral sessions from rat 107 (*left column*) and rat 108 (*right column*). See Fig. 2 captions for additional details. **a**, Left, variance explained vs. manifold dimensionality, for MIND and PCA. Right, intrinsic dimensionality score vs. manifold dimensionality. **b**, Error in decoding position from manifold coordinates, for different choices of manifold dimensionality, and from activity vectors containing the original firing rates. Error bars represent s.d. over cross-validation folds. **c**, Representation of behavior along the first two coordinates (ordered by variance) of the optimal manifold (as estimated in a). Each datapoint is colored by the values of the behavioral variables at the corresponding timepoint.

**Extended Data Figure 2:**
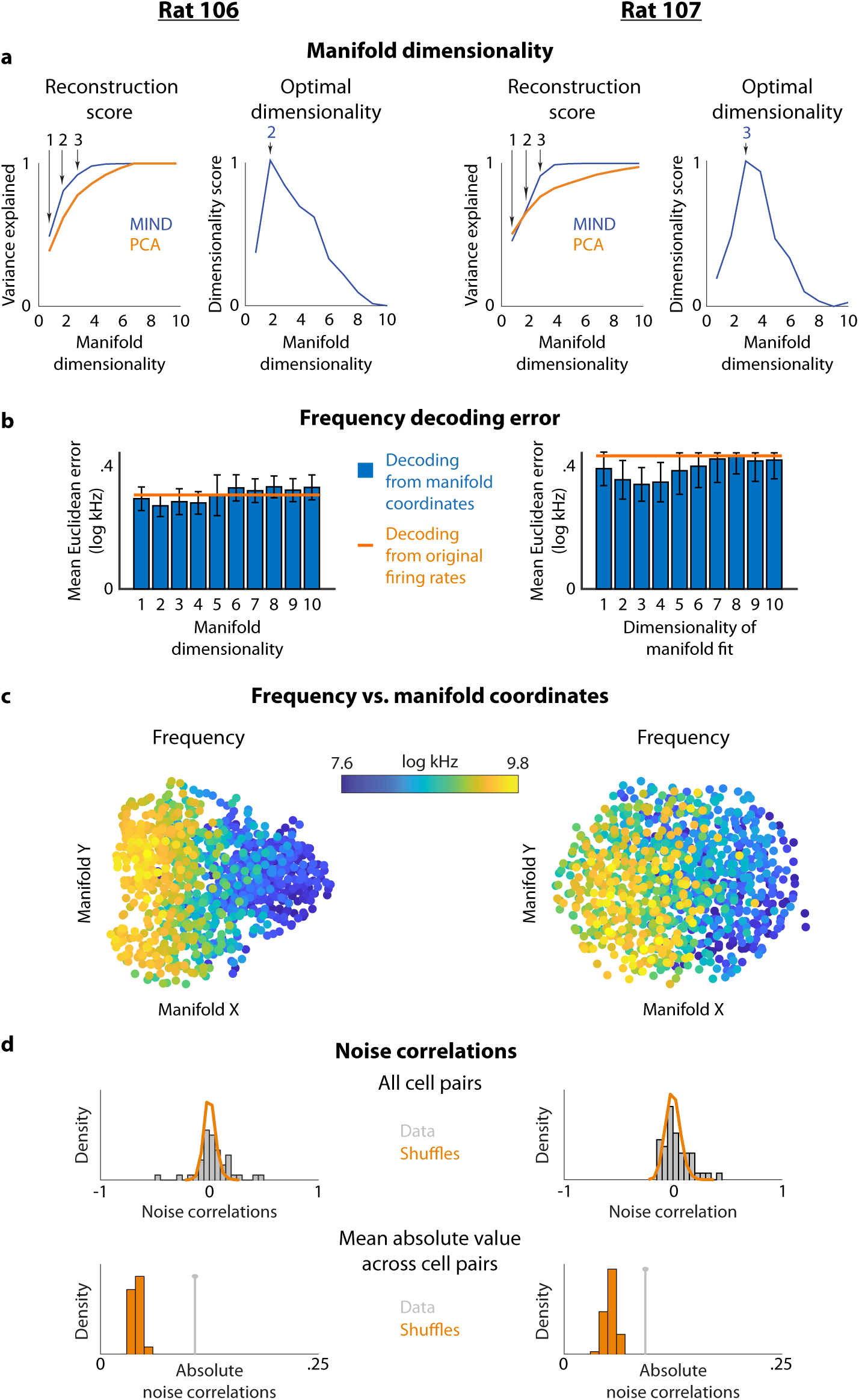
Properties of SMT manifolds for behavioral sessions from rat 106 (*left column*) and rat 107 (*right column*). See Fig. 2 captions for additional details. **a**, Left, variance explained vs. manifold dimensionality, for MIND and PCA. Right, intrinsic dimensionality score vs. manifold dimensionality. **b**, Error in decoding sound frequency from manifold coordinates, for different choices of manifold dimensionality, and from activity vectors containing the original firing rates. Error bars represent s.d. over cross-validation folds. **c**, Representation of behavior along the first two coordinates (ordered by variance) of the optimal manifold (as estimated in a). Each datapoint is colored by the values of the behavioral variables at the corresponding timepoint. **d**, Noise correlations, as in Fig. 3f. *Top:* Distribution of noise correlations across cell pairs. Shuffled datasets preserve behavior-related correlations but destroy noise correlations (see Methods). *Bottom:* Mean absolute value of noise correlations across cell pairs, compared to the same shuffles as above.

**Extended Data Figure 3:**
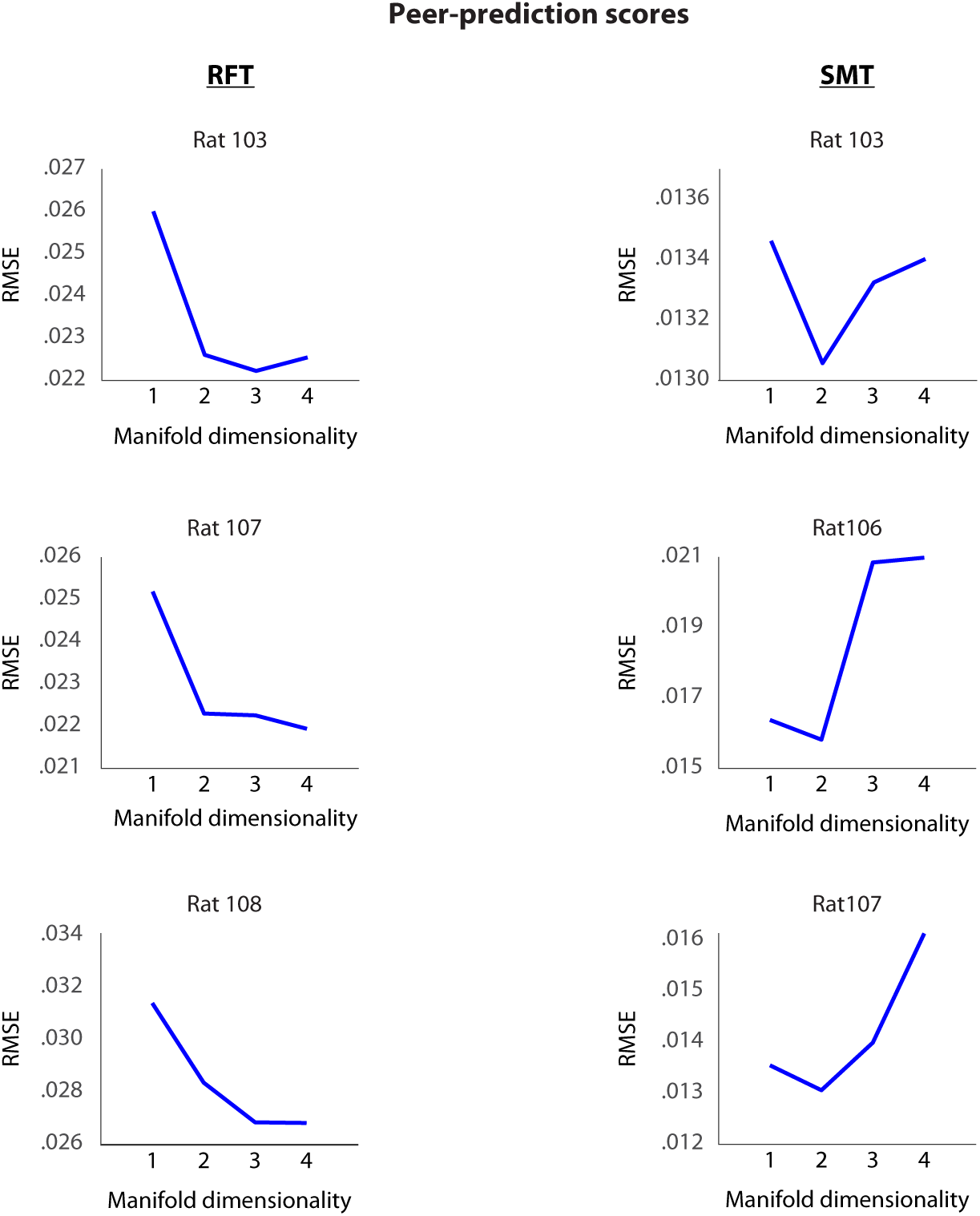
Cross-validated, leave-one-cell-out “peer-prediction scores” for the RFT (*left column*) and SMT (*right column*), as a function of manifold dimensionality. In these analyses, each cell is held out in turn, and its activity is predicted from a manifold fit to the remaining cells; the score is defined as the root mean squared error (RMSE) between the true and predicted activity, averaged over all choices of the held-out cell (see Methods for a full description). Prediction scores saturated or attained their optimum at low dimensionalities.

**Extended Data Figure 4:**
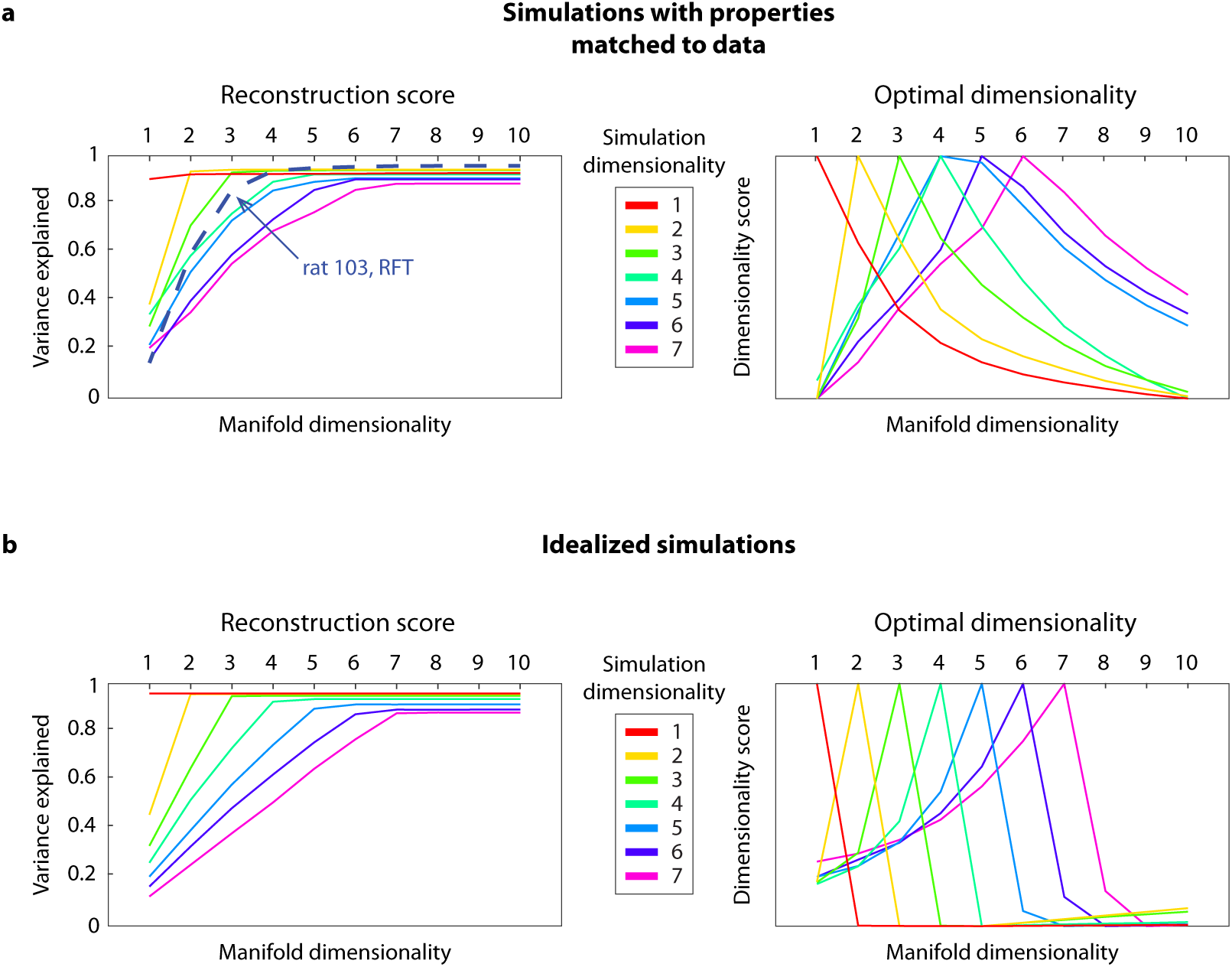
We validated our intrinsic dimensionality estimator (defined by taking the peak of the intrinsic dimensionality score) on simulated datasets of prescribed dimensionality. Simulations were modeled as place cells for a rat foraging in an arena which had the shape of a high-dimensional ball, with constant radius in all directions (see Methods). Since our estimator returns the dimensonality *d* such that the data most resembles a uniform d-dimensional ball, it should estimate the correct dimensionality on the simulations. **a**, Simulations with number of cells, number of timepoints, and firing properties of each cell matched to the RFT dataset for rat 103. *Left,* Reconstruction score. *Right,* Intrinsic dimensionality score. Undersampling does not result in an incorrect dimensionality estimate until *d* =5. At this point, our analysis requires more samples (e.g. more cells, or better exploration of the manifold) to distinguish between the manifold which generated the data, and a manifold of lower dimensionality which fits the data well. Alternatively, a correct estimate can be obtained by imposing stronger constraints on the reconstruction itself, by adjusting the hyperparameters of the MIND algorithm. **b.** The same analyses for more highly-sampled simulations (128 cells, same number of timepoints), to validate the implementation of the estimator.

**Extended Data Figure 5:**
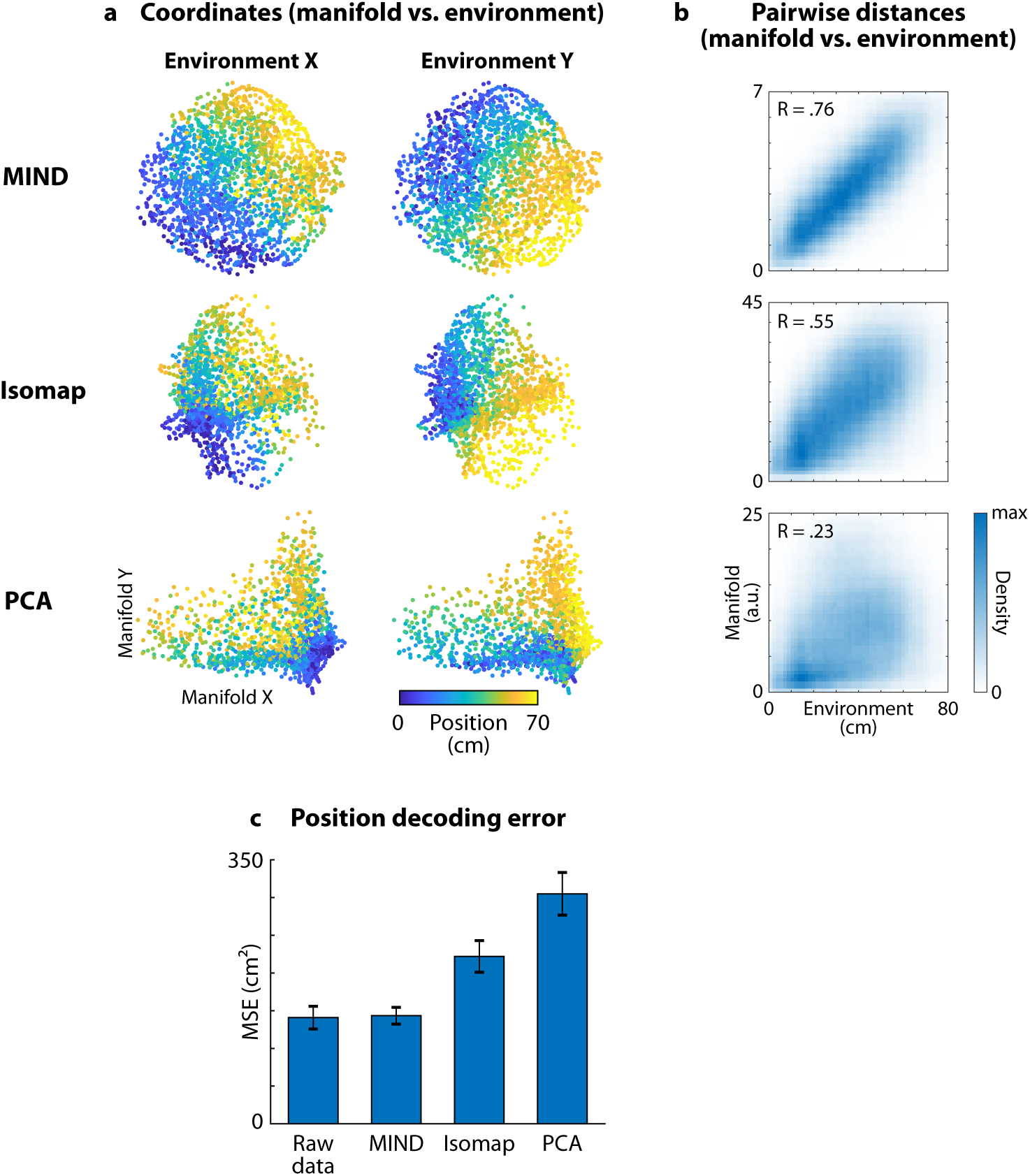
Comparison of MIND, Isomap, and PCA for capturing behavior in the RFT. Shown for rat 103. **a**, 2d manifolds fit to population activity using each method. Each point corresponds to a timepoint in the dataset. Location represents coordinates of the network on the manifold. Color represents position of the rat along each dimension of the environment. Geodesic distances for Isomap were computed using a *k* nearest neighbors graph with *k* =10 (chosen to represent position as well as possible). **b**, Geometric similarity between the 2d manifold and the environment. Quantified for each method by comparing distances between pairs of states on the manifold vs. corresponding positions in the environment. Plots show estimated joint probability density and Pearson correlation for 2 × 10^6^ pairs of points. **c**, Decoding error when predicting rat position from the high-dimensional raw data or 2d manifold coordinates using Gaussian process regression. Bar heights represent the mean decoding error (estimated using 10 fold cross-validation over temporally contiguous blocks of data). Error bars represent SEM over cross-validation folds.

**Extended Data Figure 6:**
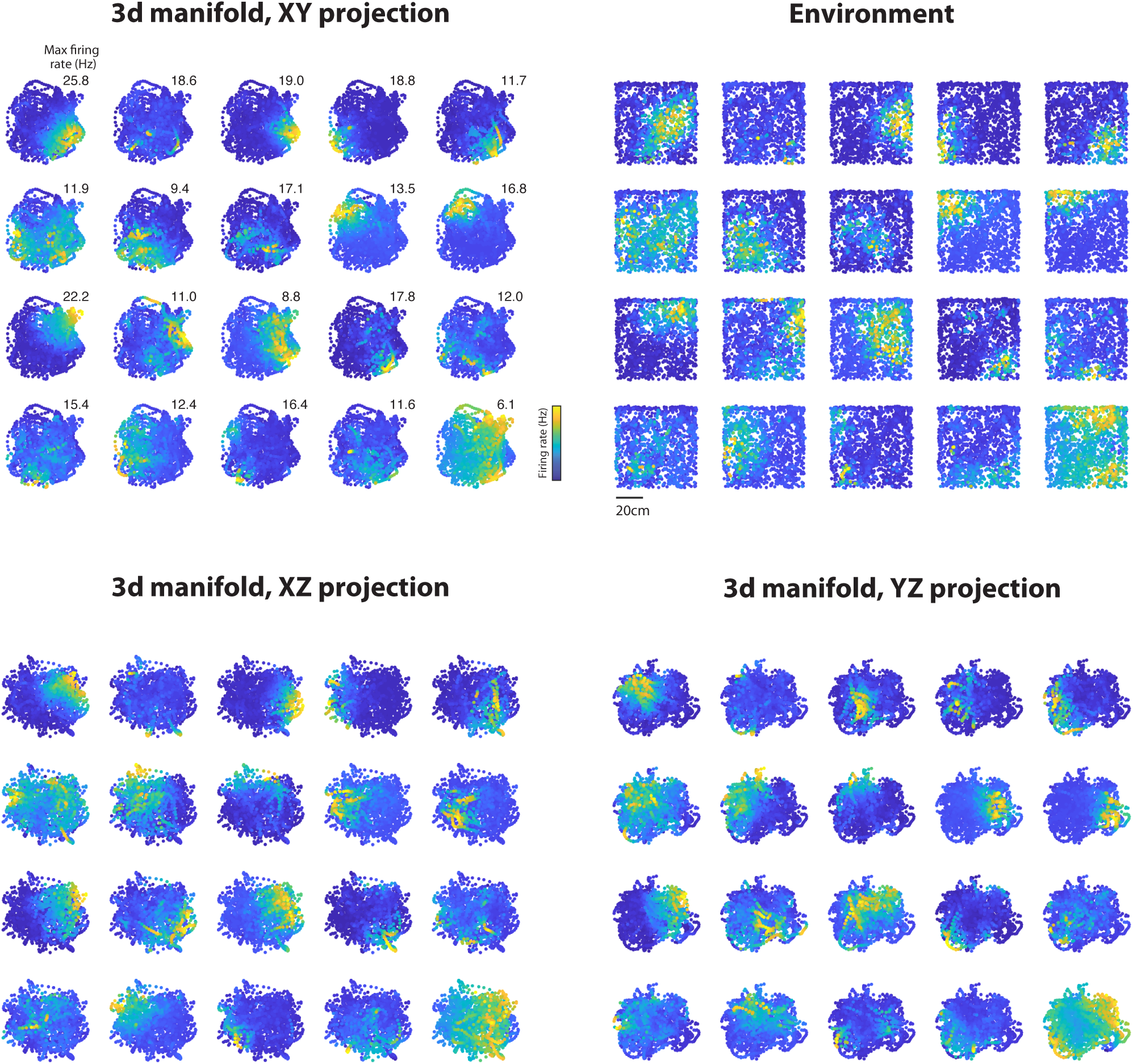
Manifold firing fields on the 3d RFT manifold. Shown for rat 103. Each field depicts cell activity vs. two of the 3d manifold coordinates. Behavioral firing fields, which depict cell activity vs. position in the environment, are also shown. Fields are shown for the twenty cells with the highest maximum firing rate across the behavioral session.

**Extended Data Figure 7:**
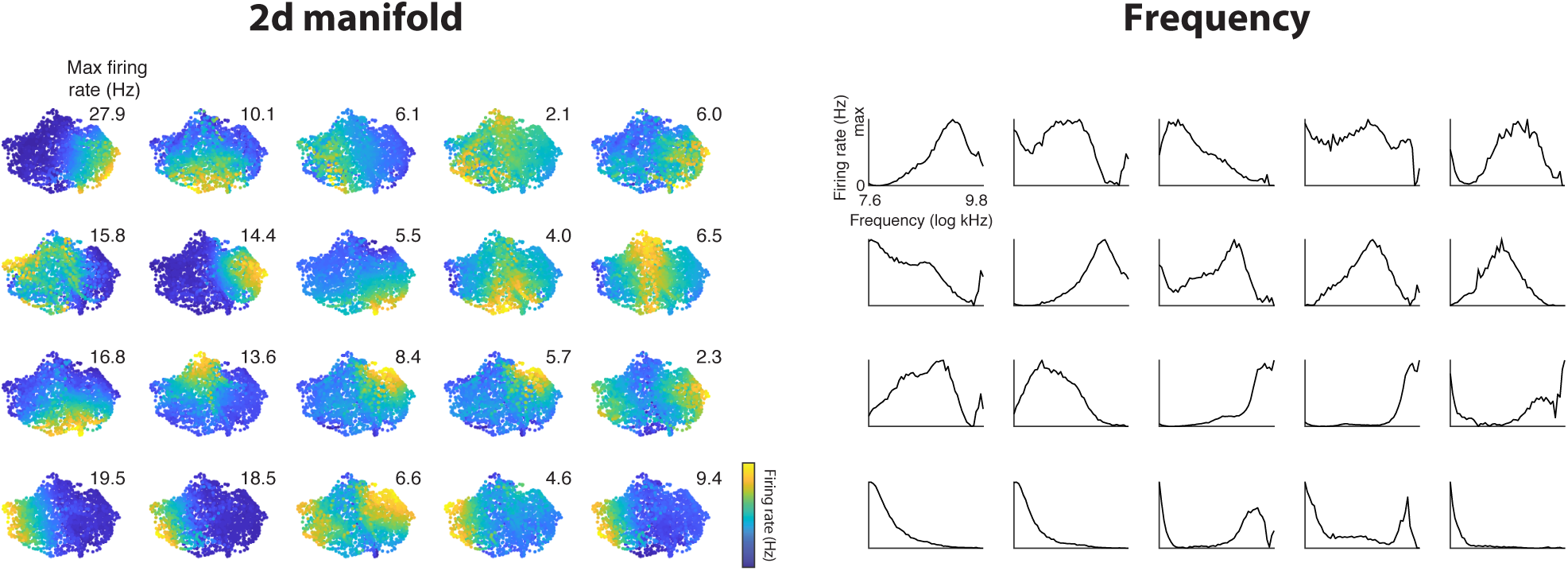
Manifold firing fields on the 2d SMT manifold. Shown for rat 103. Each field depicts cell activity vs. manifold coordinates. Behavioral firing fields, which depict cell activity vs. sound frequency, are also shown. Fields are shown for the twenty cells with the highest maximum firing rate across the behavioral session.

**Extended Data Figure 8:**
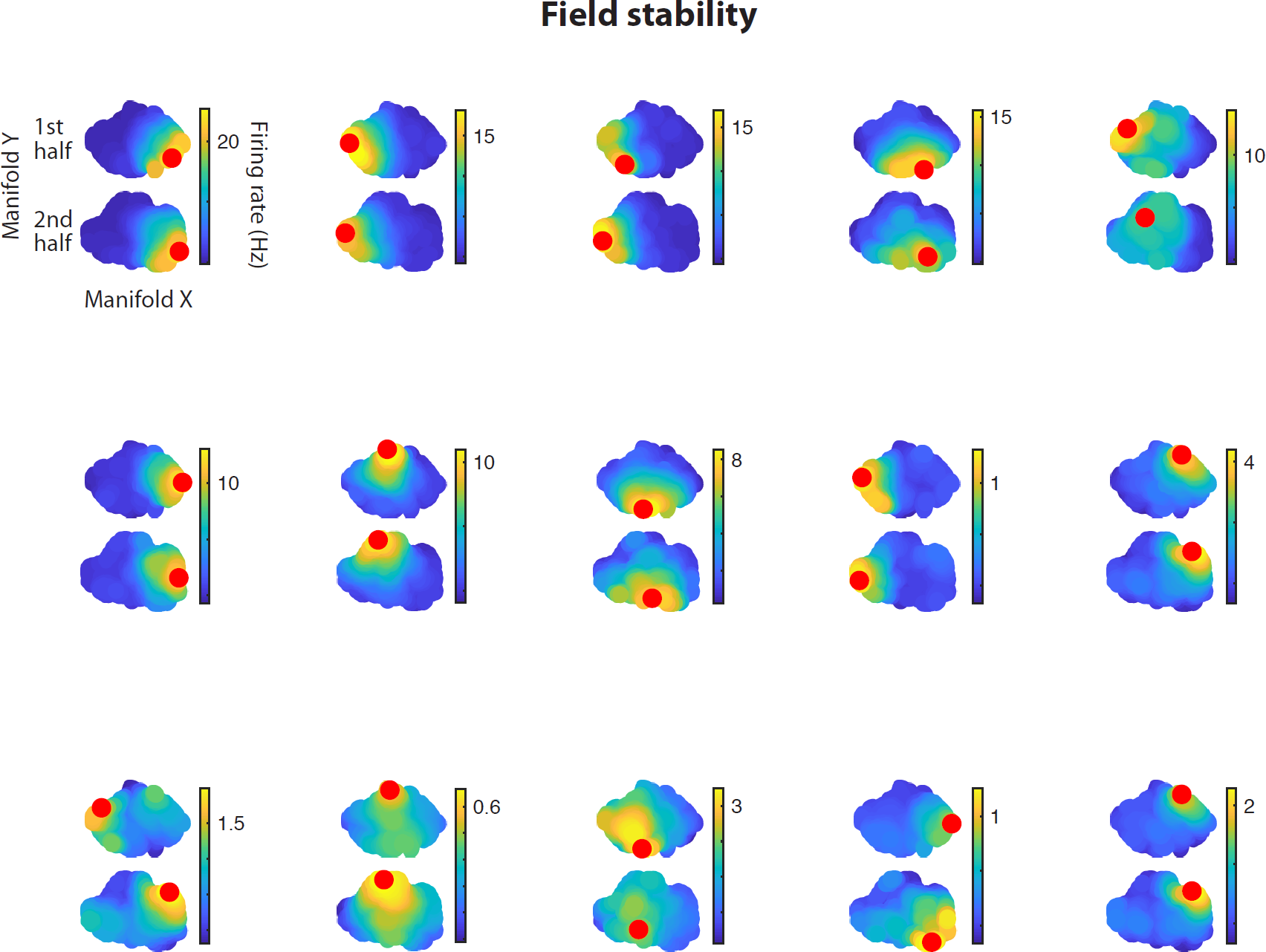
Stability of manifold firing fields in the SMT. Shown for rat 103. Each panel shows firing fields for one cell, for manifolds fit separately to the first and second half of the behavioral session (top and bottom, respectively). The center of mass of each firing field is marked in red. To align the two manifolds, each was rotated separately so that its first principle component aligned with the x axis, and the second manifold was reflected along the *x* and *y* axes if necessary to maximize field alignment between the manifolds (as measured by the Pearson’s correlation coefficient between center-of-mass vectors for each manifold, consisting of the *x* or *y* coordinates of the center of mass of each field). Fields are shown for the 15 cells with the highest maximum firing rate across the behavioral session.

**Extended Data Figure 9:**
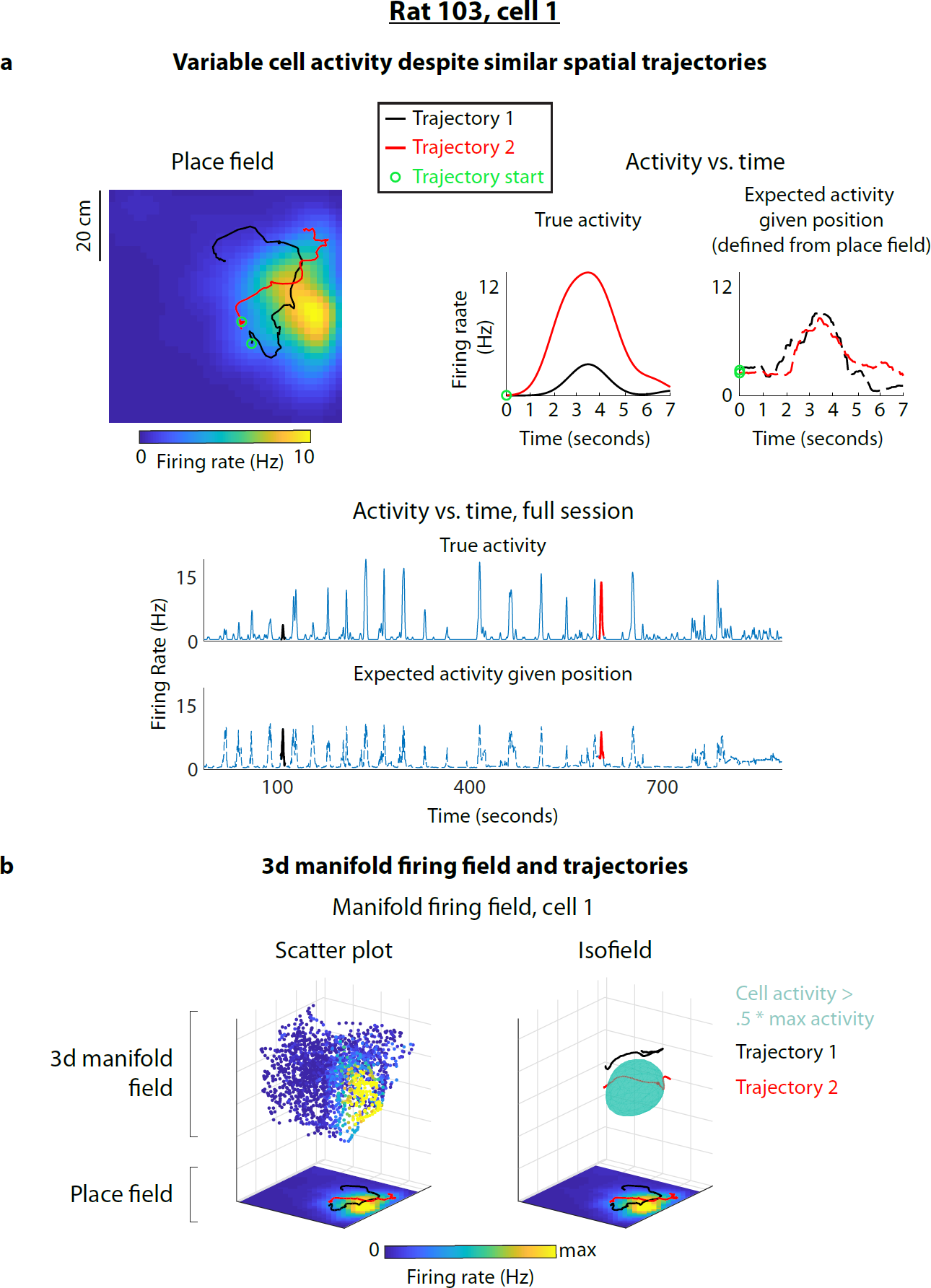
Example of a missed field on the 3d RFT manifold. Shown for rat 103. A “missed” place field occurs when the rat runs through a cell’s firing field in the environment, but the cell fails to fire (or fires an unusually low number of spikes). The missed field is explained by considering the network’s trajectory on the 3d manifold rather than the rat’s trajectory in the 2d environment. As the rat moves through the place field, the 3d neural trajectory dodges around the cell’s manifold firing field along the 3rd dimension, such that the cell does not fire. **a**, *Left,* Example of two spatial trajectory segments, each passing through the place field. The expected firing rate given position (*far right*) shows the amplitude of the place field along each spatial trajectory; the trajectories sample the place field nearly equally. However, the cell has a missed field on Trajectory 1, where the true firing rate is much lower than Trajectory 2 (*center*). Thus, the cell’s 2d place tuning cannot account for the variable neural activity. *Bottom,* For reference: the true activity and expected activity given position, over the entire session, for cell 1. **b,** *Left,* The 3d manifold firing field of cell 1. The intrinsic manifold coordinates of a representative set of datapoints are plotted, outlining the manifold. Points are colored by the firing rate of cell 1 at the corresponding timepoints. The manifold is aligned to the environment (below) by scaling, and then rotating the first two coordinates. As visualized, the place field is approximately a 2d projection of the 3d manifold field. *Right,* The 3d manifold firing field is visualized as an “isofield;” the green isosurface is defined as the level set of cell 1’s firing rate, with level equal to half the maximum amplitude. Network trajectories are plotted in the intrinsic 3d coordinate system. When projected onto the environment (below), both trajectories overlap the place field. In 3d, Trajectory 1 dodges around the manifold firing field along the 3rd dimension, whereas Trajectory 2 passes through its center. Thus, the interaction between network trajectories and firing fields on the 3d manifold accounts for the variable neural activity.

**Extended Data Figure 10:**
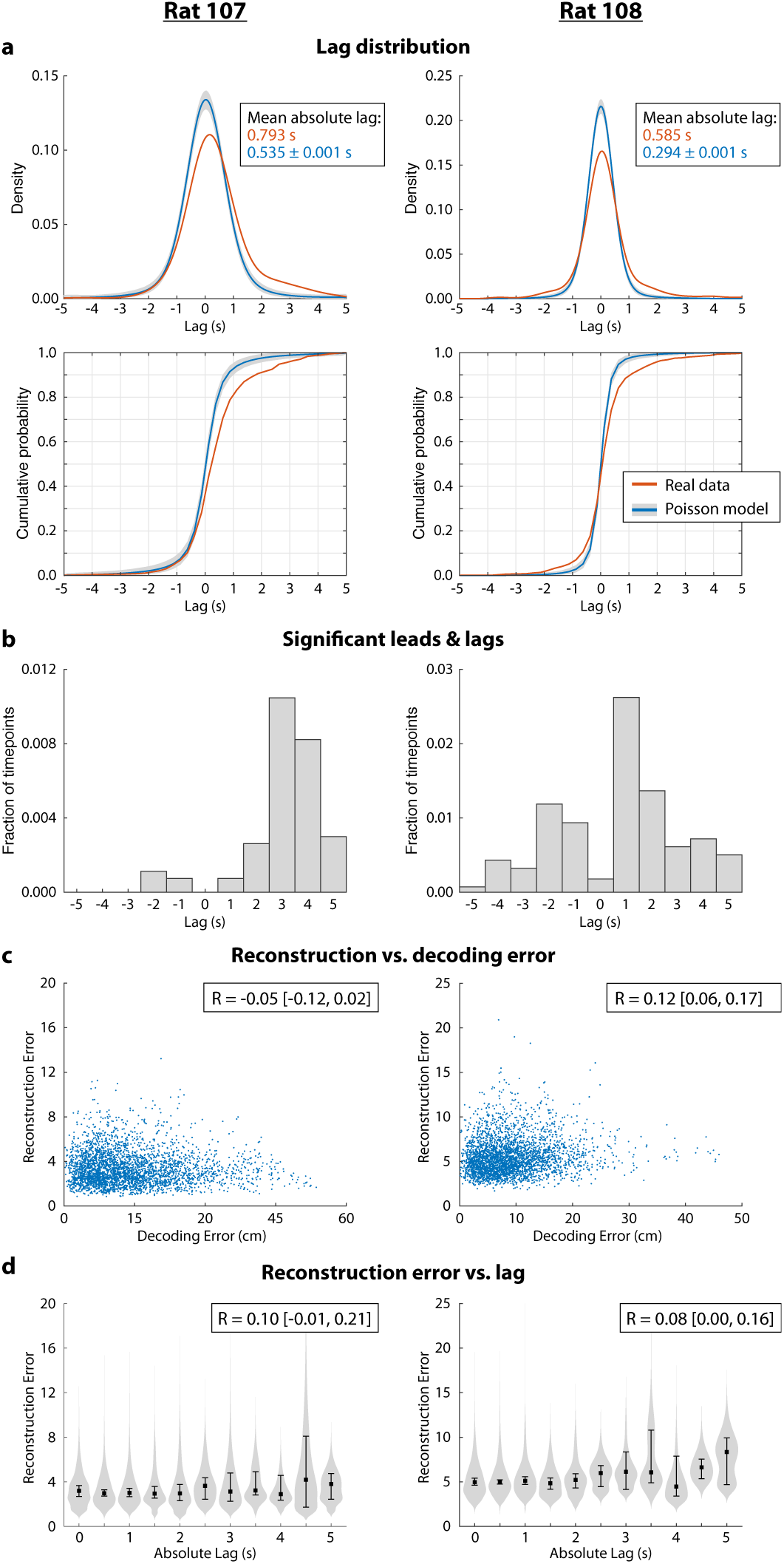
Analysis of deviations from encoding current position in the RFT, for rats 107 (left column) and 108 (right column), in the behavioral sessions from Extended Data Fig. 1. **a**, Estimated time lag between the predicted and actual position. Top: probability density function of lag aggregated across time (kernel density estimate). *Bottom*: Corresponding cumulative distribution function (empirical CDF). The gray region is a pointwise 95% confidence band and the blue curve is the average across synthetic Poisson datasets. **b**, The relative frequency of statistically significant timepoints with each lag, expressed as a fraction of the total number of timepoints included in the analysis. **c**, Manifold reconstruction error vs. decoding error. Pearson correlation is shown with 95% bootstrap confidence intervals. **d**, Manifold reconstruction error vs. absolute lag. Points and error bars show the median and 95% bootstrap confidence intervals. Gray regions (violin plot) show the conditional density of reconstruction error given absolute lag. Pearson correlation is shown with 95% bootstrap confidence intervals.

**Extended Data Figure 11:**
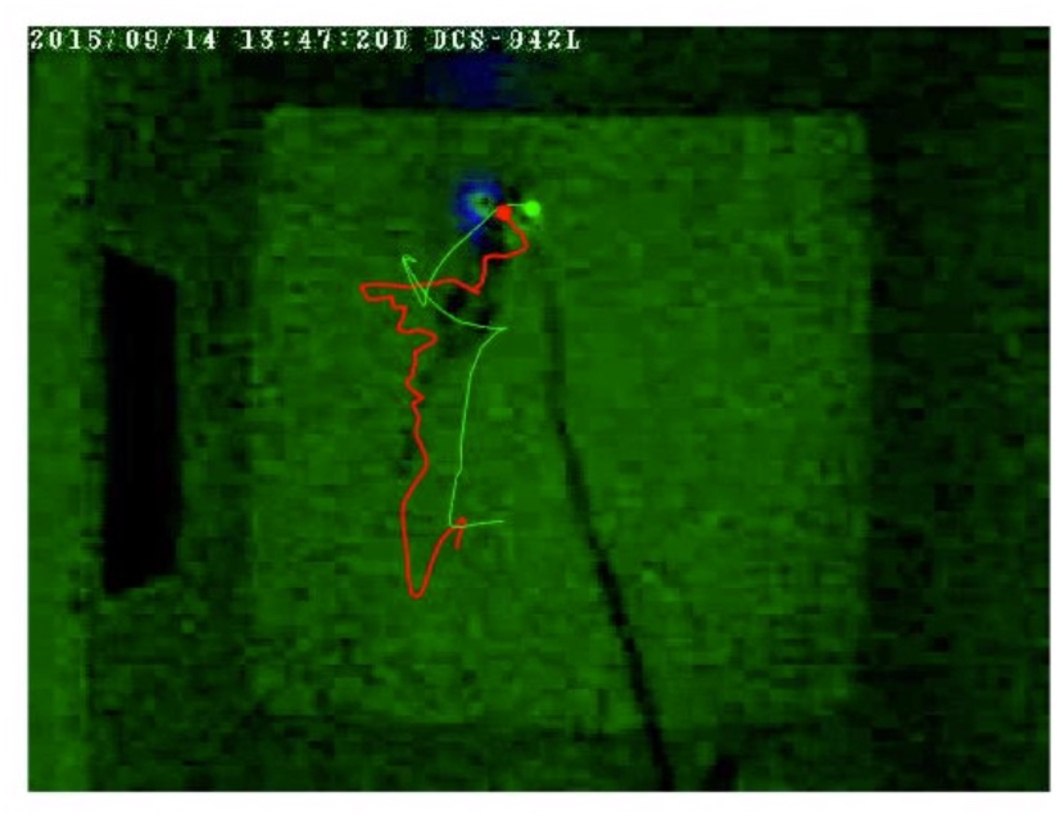
Legend for trajectory video. Behavioral video excerpt for an RFT session (rat 103). Markers overlaid on each frame show the rat’s true position (red) and predicted position (green), given the network’s location on the manifold. The past 8 s of the real and decoded behavioral trajectories are shown as red and green lines, respectively. The following annotations indicate deviations from encoding the current position, as described in Fig. 4. During detours, the green decoded trajectory becomes white. During leads and lags, yellow and blue markers appear on both the real and decoded trajectories, indicating the segments where the decoded trajectory is leading or lagging the real trajectory, respectively. Leads and lags were detected by statistical comparison to the Poisson null model, as in Fig. 4c. See Methods for lead, lag, and detour detection.

## Methods

### 1 Manifold discovery

#### 1.1 Inputs

Input consists of population activity over time, and exogenous behavioral variables are not used. Activity for a population of N neurons is given as a time series of network states 𝒳 = {*x*_1_,*…,x*_T_} where each state *x*_*t*_ ∈ ℝ^*N*^ is an *N*-dimensional vector in a continuous state space, with each dimension representing the activity of a neuron at time *t*.

#### 1.2 Modeling state transitions using PPCA forests

A probabilistic model of transitions between states is then learned. Let *X*_*t*_ be a continuous random vector representing the state at time *t*, and let *X*_*t*+1_ represent its “successor state,” the state one timestep later. The conditional distribution over successor states given current state *X*_*t*_ is *p*(*X*_*t*+1_ | *X*_*t*_), and is assumed not to depend on time. Therefore, for states *x, x*′ ∈ ℝ^*N*^, *p*(*X*_*t*+1_ = *x*′ | *X*_*t*_ = *x*) is the same for all *t*, and we will write this as as *p*(*x*′| *x*) for convenience. We refer to this as the transition probability density function.

The transition probability density function is estimated using a novel algorithm that employs decision trees to adaptively partition state space into local neighborhoods, and probabilistic PCA to model the distribution over successor states for each neighborhood. Distributions from a large set of randomized trees are then combined to give an ensemble model with better performance than any single tree. This algorithm, which we call ‘PPCA forests’, combines ideas from probabilistic PCA,^1^ random projection trees,^2,3^ and random forests.^4^

##### 1.2.1 Modeling state transitions using decision trees

Decision trees are used to model the conditional distribution over successor states, given the current state. In this approach, state space is partitioned into local neighborhoods, with a local distribution over successor states for current states that lie in each neighborhood. This strategy is similar to tree-based regression methods,^5^ but operates in an unsupervised manner (there are no exogenous target variables), and produces probability distributions as outputs rather than point estimates.

A decision tree represents a hierarchical partition of state space, where each node of the tree corresponds to a local region 𝒜_*i*_ of state space (where i is the node index). The root node represents the entire space. Each internal node has a direction parameter v and threshold parameter q that define a hyperplane. The hyperplane splits 𝒜_*i*_ into subregions 𝒜_*L*(*i*)_ and 𝒜_*R*(*i*)_, corresponding to the left and right child nodes *L*(*i*) and *R*(*i*) in the tree.

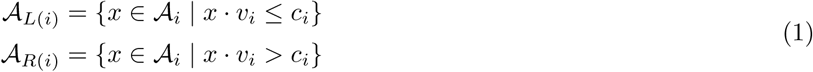

Leaf nodes, which have no children, represent the finest partition of state space into disjoint subregions. The index of the leaf node containing state *x* can be found by traversing the decision tree. Starting from the root node, *x* is recursively assigned to the left or right child node according to (1), until hitting a leaf node.

Each leaf node has an associated Gaussian distribution with mean *μ*_*i*_ and covariance matrix *C*_*i*_, which models the distribution over successor states, given that the current state falls within 𝒜_*i*_. As a whole, the decision tree implements an estimated transition probability density function *f* (*x*′ | *x*), which assigns a probability density to possible successor state *x*′, given current state *x*. f is computed by traversing the tree to find the leaf node containing *x*. In terms of this leaf node, the estimated transition probability density function is given by

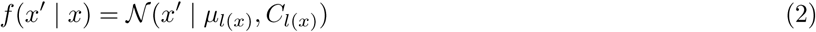

where *l*(*x*) is the index of the leaf node containing *x*, and 𝒩(*x* | *μ, C*) denotes a Gaussian density function with mean *μ* and covariance matrix *C*.

##### 1.2.2 Fitting decision trees

A decision tree is learned by recursively partitioning state space into progressively finer neighborhoods. A tree begins as a single leaf node (the root) representing the entire state space. At each step, a leaf node is split to create two child nodes, and a local model is fit to the successors of states in the dataset that fall within each child region. Splitting parameters and local models are jointly optimized, such that local neighborhoods adapt to the data. However, random constraints are imposed, which serve to decorrelate the trees and increase the accuracy of the final ensemble model. The training procedure can be seen as a randomized, greedy algorithm for approximately maximizing the likelihood of each tree.

###### Learning local distributions with probabilistic PCA

The distribution over successor states for each neighborhood is modeled as a Gaussian, whose mean and covariance matrix must be estimated from the data. The trajectory through state space is believed to lie near a low-dimensional, nonlinear manifold. This implies that the distribution over states in a sufficiently small region should be concentrated near the local tangent plane of the manifold with small out-of-plane deviations due to noise and the curvature of the manifold. Variance should therefore be low along directions orthogonal to the tangent plane. To exploit this low-dimensional structure, probabilistic PCA^1^ (PPCA) is used to estimate the distribution over successor states. PPCA provides regularization by imposing low-dimensional constraints on the covariance matrix, thereby reducing overfitting.

Let 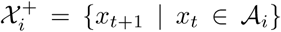 denote the set containing the successors of states in the dataset that fall within neighborhood 𝒜_*i*_. The mean *μ*_*i*_ and covariance matrix *C*_*i*_ are estimated from 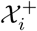 using maximum likelihood PPCA, as described below (Section 1.7).

###### Splitting rules

Splitting a node creates two child nodes, each with a local PPCA model. Splitting parameters determine the data assigned to each child node, and are chosen to maximize the likelihood of the resulting PPCA models.

Parameters for splitting node *i* consist of a direction and threshold that define a hyperplane separating the local neighborhood 𝒜_*i*_ into two subregions (1). Suppose that 𝒜_*i*_ were split along direction *v* with threshold *c*. Let sets 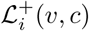 and 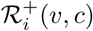 contain the successors of states in the dataset that fall within the resulting left and right subregions.

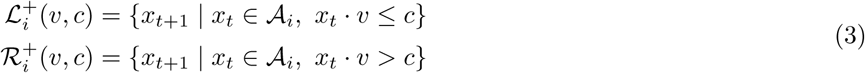

PPCA models are fit to each of these sets (see Section 1.7). Let Γ_*ML*_(•) denote the log likelihood of the maximum likelihood PPCA model for a given set of points, computed as in (15). The splitting direction *v*_*i*_ and threshold *c*_*i*_ are chosen as:

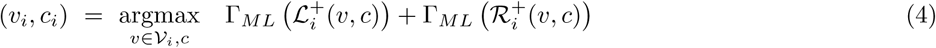

The splitting direction is constrained to 𝒱_*i*_, a set of *n*_*dir*_ < *N* random directions sampled from the uniform distribution on the unit sphere. This constraint introduces randomness into the fitting procedure, which decorrelates trees and plays an important role when multiple trees are combined into an ensemble model (Section 1.2.3). The splitting threshold is constrained such that no fewer than *n*_*leaf*_ data points are assigned to either subregion. Choice of the hyperparameters *n*_*dir*_ and *n*_*leaf*_ is discussed below.

The optimization problem (4) can be solved by iterating through directions and a finite number of thresholds that uniquely partition the data along each direction. PPCA models are fit for each candidate direction and threshold, and the direction, threshold, and PPCA parameters that maximize the log likelihood are retained. We describe ways to accelerate the optimization procedure in Section 1.6.2.

In general machine learning applications (e.g. classification, regression, and nearest neighbor search), decision tree nodes are commonly split along single dimensions. This corresponds to splitting with axis-parallel hyperplanes, producing hyperrectangular local regions. Here, we perform splits along randomized, oblique directions, producing local regions that are convex polytopes. This strategy allows trees to adapt more efficiently to data distributed along low-dimensional manifolds, but embedded in spaces with high ambient dimension.^2,3^

###### Stopping rules

determine when the recursive splitting procedure terminates, producing a final leaf node. A node will not be split if it contains fewer than 2*n*_*leaf*_ data points. Together with constraints on the splitting threshold (4), this ensures that every leaf node in the final model contains at least *n*_*leaf*_ data points. Similar to regression trees and random forests,^4,5^ decreasing *n*_*leaf*_ produces a larger number of smaller neighborhoods, which increases flexibility of the overall model and reduces bias. However, it also leaves fewer data points available for fitting local models, which increases variance. As in random forests,^4^ this cost is offset by combining the outputs of many trees, which reduces variance, thereby allowing more flexible trees (see Section 1.2.3). A general guideline is that *n*_*leaf*_ must be large enough to allow the estimation of a local PPCA model. *n*_*leaf*_ can be selected using cross-validation. In our analyses, we used *n*_*leaf*_ = 40, which was chosen in an *ad hoc* manner, and not optimized using the data.

##### 1.2.3 Combining trees into a forest

Individual trees are combined into a forest, which is an ensemble model of transitions between network states. Each tree represents a partition of state space into local neighborhoods, with a transition model for each neighborhood. Because node splitting directions are chosen from a random set (4), the neighborhoods and local models are different for each tree. This randomization serves to decorrelate trees. As with random forests^4^ and bagging,^5^ combining the outputs of a diverse set of models reduces variance, yielding an ensemble model with greater performance than any individual model.

Let {*f*_1_,…,*f*_s_} denote the models for *S* trees, each specifying a transition probability density function as in (2). Given current state *x*, the combined transition probability density function is:

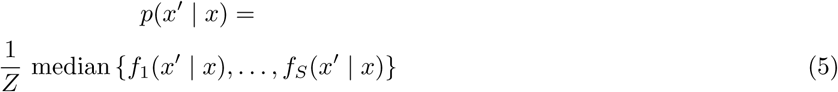

The sample median of the individual model outputs is used for robustness. *Z* is a normalizing constant such that the distribution integrates to one. In practice, it is not necessary to compute *Z*, as further analyses only require values proportional to the probability density.

The degree of dependence between trees is controlled by the hyperparameter *n*_*dir*_, which specifies the number of random directions to consider when splitting a node (4). *n*_*dir*_ plays a similar role as the hyperparameter in random forests that governs the number of input features to consider on each split.^4,5^ Analogously, increasing *n*_*dir*_ allows individual trees to fit better by choosing more optimal splitting directions, which decreases bias. However, it also increases correlation between trees, which increases variance—intuitively, the benefit of ensembling is reduced as the individual models become less diverse. The optimal setting of *n*_*dir*_ involves a tradeoff between these opposing effects, and can be estimated using cross-validation. In our analyses, we used *n*_*dir*_ = 2, which was chosen in an *ad hoc* manner, and not optimized using the data.

Randomized trees are fit independently, and the model combination procedure itself doesn’t involve any fitting. Thus, as in random forests, there is no risk of overfitting by increasing the number of trees *S*.^4,5^ Rather, increasing *S* serves to decrease variance, thereby increasing accuracy of the ensemble. Therefore, S should be set as high as computationally feasible. However, performance eventually asymptotes, so there is little additional benefit to increasing *S* beyond a certain point. In our analyses, we used *S* = 100.

#### 1.3 Measuring distance between network states

The model of state transitions learned in Section 1.2 was used to construct a distance measure between states. Distances between points on a low-dimensional manifold can be used to estimate geometric and topological properties of the manifold, and to map points to low-dimensional, intrinsic coordinates on the manifold.^6^ The choice of distance measure plays an important role in estimating these quantities from the data.

Manifold learning algorithms are often based on predefined distance measures in input space (e.g. Euclidean distance). In our application, this would correspond to comparing the firing rates of each neuron in different states. However, it is unclear *a priori* how to measure the similarity of firing rates in a way that reflects their meaning or influence on the network. Instead, we sought to learn distances from the behavior of the network itself.

We defined a novel distance measure based on transition probabilities, which captures structure in the dynamics of the system. For example, two states are assigned a small distance if the network transitions directly between them with high probability, and a large distance if the network only transitions between them by moving through many intermediate states. This notion of distance is agnostic to the nature of the states, and measures only how the network transitions between them. Our distance measure is based on arranging points on a manifold such that the transition probabilities of a random walk on the manifold match those estimated from the data.

##### 1.3.1 Local distances

The PPCA forest models transitions over the entire, continuous state space. Restricting transitions to states in the dataset gives a discrete approximation to this process, captured by stochastic matrix *P*. Let *P*_*ij*_ denote the probability of transitioning from state *x*_*i*_ to *x*_*j*_:

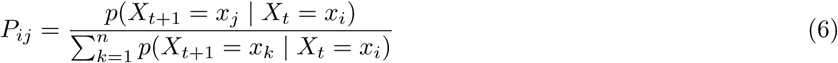

where *P*(*X*_*t*+1_ | *X*_*t*_) denotes the PPCA forest output.

Suppose each state *x*_*i*_ in the dataset has an image *y*_*i*_ on a manifold. Consider a random walk that traverses these points, acting as a discrete diffusion process on the manifold. The probability of transitioning from *y*_*i*_ to *y*_*j*_ can be approximated by an isotropic Gaussian kernel function:

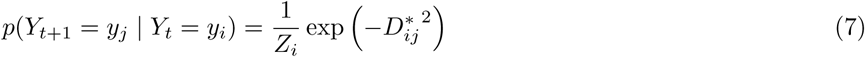

where 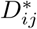 denotes local distance on the manifold from *y*_*i*_ to *y*_*j*_ and *Z*_*i*_ is a normalizing constant such that transition probabilities sum to one. For convenience, we assume *Z*_*i*_ = 1, as distances can be adjusted to compensate.

Assume transition probabilities of the random walk are equal to those estimated from the data. Setting random walk transition probabilities to *P*_*ij*_ and solving for distances gives:

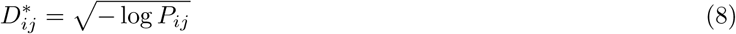

##### 1.3.2 Global distances

When calculated as in (8), *D*^*^ measures ideal *local* distances on the manifold. However, it is infinite when the estimated transition probability between states is zero, and is not symmetric unless estimated transition probabilities are symmetric. We used local distances to define a global, symmetric distance measure suitable for manifold learning.

Similar to isomap,^7^ we defined the global distance between two states as the length of the shortest path from one to the other via any intermediate, connected states. Consider a directed graph *G* whose vertices correspond to states. An edge from vertex *i* to vertex *j* exists with weight 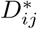 if there is nonzero probability of transitioning from state *i* to state *j*. The global distance from state *i* to state *j* is given by 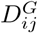, the geodesic distance on *G* from vertex *i* to vertex *j*. Geodesic distances between all pairs of states were computed using Johnson’s algorithm.^8^

Geodesic distances were then symmetrized to give the final, global distance:

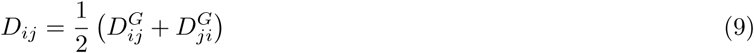

#### 1.4 Learning intrinsic manifold coordinates

Points on the manifold can be indexed by a set of low-dimensional coordinates. Intrinsic manifold coordinates were obtained for observed network states by embedding them into a vector space such that distances on the manifold were approximately preserved. Given pairwise distance matrix *D* (as estimated above), d-dimensional intrinsic coordinates {*y*_*i*_,…, *y*_*T*_} ⊂ ℝ^*d*^ were obtained for each data point by solving the following optimization problem:

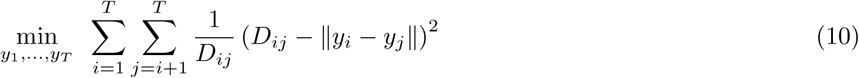

This problem corresponds to Sammon mapping^9^—a form of nonclassical, metric multidimensional scaling (MDS)^10^— using our novel distance measure that captures structure in dynamics. In (10), we seek embedding coordinates such that Euclidean distances in the embedding space best match the given distances in the least squares sense. The weight term 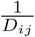 places greater emphasis on preserving distances between points that are nearby on the manifold. This choice reflects the importance of local distances, in that: 1) Local distances are directly computed from estimated transition probabilities, whereas global distances are derived from them; and 2) Local distances are of primary relevance to recovering the local topology of the manifold. However, the fact that global distances do contribute to the embedding may provide robustness over purely local embedding techniques.^6^ The dimensionality *d* is a hyperparameter, and can be selected using the method described in Section 6.

We solved (10) using the conjugate gradient method. The optimization was initialized with the *d*-dimensional classical MDS solution for distance matrix *N*, which can be cast as an eigenproblem and solved in closed form.^10^ The solution to (10) is only specified up to rigid transformations. To obtain a canonical representation, we used PCA to center and rotate the solution (without discarding any dimensions). This ensured that the resulting manifold coordinates of the data were uncorrelated, aligned with the directions of greatest variance, and ordered by decreasing variance.

#### 1.5 Mappings between state space and the manifold

##### 1.5.1 Forward mapping

The above procedure (Section 1.4) produces intrinsic coordinates on the manifold for each network state in the dataset. However, it is also useful to have an explicit mapping from state space to the manifold, so that manifold coordinates can also be obtained for out-of-sample data that are not present in the training set. We treat this as a supervised learning problem: Given states 𝒳 = {*x*_1_,…, *x*_*T*_} ⊂ ℝ^*N*^ with corresponding manifold coordinates 𝒴 = {*y*_1_,…, *y*_*T*_} ⊂ ℝ^*d*^, learn a mapping *gf* : ℝ^*N*^ → ℝ^*d*^ such that *gf* (*x*_*i*_) approximates *y*_*i*_ for all *i*.

In principle, any sufficiently powerful, nonlinear regression method with multidimensional outputs could be used for this purpose. We used a nonparametric regression method originally proposed for use with locally linear embedding (LLE).^11^ To map a new point *x* in state space onto the manifold, its *k* nearest neighbors {*η*_1_(*x*),…, *η*_*k*_(*x*)} ⊂ 𝒳 are found, as measured by Euclidean distance in state space. *x* is then expressed as a weighted sum of its neighbors. Weights *w*(*x*) are found that minimize the squared reconstruction error plus an *l*_2_ regularization term that encourages small weights. Weights are constrained to sum to one, which makes them invariant to translation of x and its neighbors (they are also invariant to rotation and scaling).

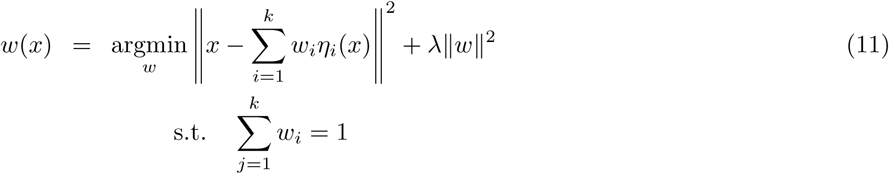

An efficiently computable, closed form solution to this optimization problem is described in.^11^ Let G denote the local Gram matrix:

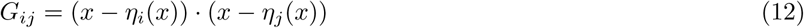

Let *I* denote the identity matrix and 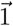 denote a vector of ones. The optimal weights can be found by solving the following equation and then rescaling the weights so that they sum to one.

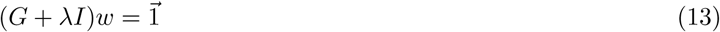

*x* is then mapped onto the manifold by applying the same weights to the known images of its neighbors on the manifold 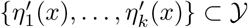.

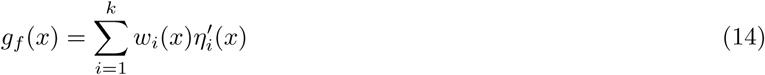

The number of neighbors *k* and the regularization strength λ are hyperparameters, and can be chosen using cross-validation on the original dataset (where coordinates in state space and on the manifold are both known). In our analyses, we used 10-fold cross-validation.

##### 1.5.2 Reverse mapping

A similar approach can be used to obtain an explicit mapping from the manifold back to state space. The reverse mapping gives the corresponding pattern of neural activity for any point on the manifold. As above, we treat this as a supervised learning problem: Given states 𝒳 = {*x*_1_,…,*x*_*T*_} ⊂ ℝ^*N*^ with corresponding manifold coordinates 𝒴 = {*y*_1_,…, *y*_*T*_} ⊂ ℝ^*d*^, learn a mapping *g*_*r*_ : ℝ^*d*^ → ℝ^*N*^ such that *g*_*r*_(*y*_*i*_) approximates *x*_*i*_ for all *i*. The nonparametric regression method described above can be used here, by reversing the roles of 𝒳 and 𝒴: 1) Given a new point *y* on the manifold, find its *k* nearest neighbors in the training set. 2) Find the optimal weights that approximate *y* as a linear combination of its neighbors. 3) Obtain *g*_*r*_(*y*) by applying the same weights to the known coordinates of the neighbors in state space. As above, the hyperparameters k and A can be selected using cross-validation. The optimal hyperparameter values for the forward and reverse mappings will generally differ.

#### 1.6 Computational considerations

##### 1.6.1 Preprocessing with PCA

The data can be preprocessed using principal component analysis (PCA) to reduce its dimensionality, thereby speeding up further computations. In this case, the network states described above (Section 1.1) represent the component scores at each timepoint, rather than the raw activities of neurons. The ambient dimensionality *N* of state space then corresponds to the number of components. We used PCA preprocessing in our analyses, and chose *N* to preserve at least 95% of the variance of the raw neural activity. This gives a lower-dimensional linear subspace (in which a nonlinear manifold of even lower intrinsic dimensionality is embedded), without discarding much information.

Dimensionality reduction decreases the time needed to estimate transition probabilities, and to evaluate mappings between state space and the manifold. For example, fitting the PPCA forest requires multiple eigenvalue decompositions of covariance matrices, each requiring *O*(*N*^3^) time. Evaluating the nonparametric regression models described in Section 1.5 involves pairwise distance computations whose runtime scales linearly with *N*. In some cases, dimensionality reduction may give the additional benefit of denoising the data.

Note that, when PCA preprocessing is used, the mappings described above (Section 1.5) relate component scores to manifold coordinates. In this case, raw neural activity can easily be related to manifold coordinates by transforming raw activity to component scores (and vice versa) using the simple forward and reverse mappings provided by PCA.

##### 1.6.2 Accelerating PPCA forest training

Training the PPCA forest requires fitting multiple local PPCA models. A node is split at each step of fitting an individual decision tree, which requires selecting a splitting direction and threshold from a set of candidate values. The direction and threshold are chosen to maximize the likelihood of the PPCA models for the child nodes created by the split. This requires fitting two PPCA models for each possible choice of splitting direction and threshold.

Given a splitting direction, an exhaustive search over thresholds may be computationally prohibitive. To speed up training, a smaller set of thresholds can be tested instead. For example, we implemented the following procedure. Given a candidate splitting direction, the best threshold is selected from a small number of candidates (e.g. 10 or 20) that divide the data into even quantiles along the given direction. Golden section search is then used to optimize the threshold, starting from this initial value.

Alternatively, to speed up training even further, we use the following approximation procedure. Given a splitting direction, the threshold is chosen such that the mean successor state for each child node best predicts the actual successor states, as measured by the squared error. This is equivalent to maximizing the likelihood, assuming that the distribution over successor states for each child node is an isotropic Gaussian distribution. This optimization problem does not require estimating or decomposing covariance matrices, and can be solved efficiently using running sums to update the error for each choice of threshold, rather than recomputing it from scratch.^12^ Once the threshold has been chosen, full PPCA models are then fit to the successor states for each child node. This procedure is repeated for each candidate splitting direction, and the final direction and threshold are chosen to maximize the likelihood of the PPCA models. Therefore, only a single PPCA fit per direction is required.

##### 1.6.3 Landmark approach

Once transition probabilities have been estimated, learning the manifold structure as described above involves computations on distances between all pairs of network states in the dataset. For *T* timepoints, this requires *O*(*T*^2^) memory. Computing geodesic distances using Johnson’s algorithm has time complexity *O*(*T* log*T* + *T*|*E*|) (where |*E*| denotes the number of pairs of network states with nonzero transition probability). Finding manifold coordinates using Sammon mapping has time complexity *O*(*iT*^2^) (where i denotes the number of iterations). These requirements can become infeasible for datasets with many timepoints.

To enable manifold discovery with large datasets, we employ a ‘landmark approach’ similar to that described in^13^. The PPCA forest is fit using the full dataset. Thereafter, distances are computed between a representative subset of states called ‘landmark points’, which are used to learn the manifold structure. Thus, computational complexity depends only on the number of landmark points *T*′ ≪ *T*. In our analyses, we used *T*′ = 2000, allowing distance computations and manifold learning in just a few minutes on standard hardware.

We selected landmark points as the medoids obtained via k-medoids clustering^5^ of all states in the dataset (using Euclidean distances between states). For very large datasets, landmark points could instead be chosen using an efficient, greedy algorithm described in.^13^ Using the landmark approach, intrinsic manifold coordinates are obtained for each landmark point. To obtain full trajectories on the manifold, all states can then be mapped onto the manifold using the forward mapping described above (Section 1.5).

#### 1.7 Probabilistic PCA

Probabilistic PCA (PPCA)^1^ is a probabilistic variant of classical PCA, formulated as a Gaussian latent variable model. We use it as a means to fit a Gaussian distribution with low-dimensional constraints on the covariance matrix—that is, the variance along many directions should be small.

Let *Ƶ* = {*z*_*i*_,….,*z*_*T*_} ⊂ ℝ^*N*^ denote a set of generic data points. Given a specified dimensionality *q*, PPCA models the data with a Gaussian distribution with mean *μ* and covariance matrix *C* = *WW*^*T*^ + *σ*^2^*I*, where *W* is a *N* × *q* matrix, *I* is the identity matrix, and *σ*^2^ is the noise variance. This expression for *C* constrains the variance along each of the smallest *N* − *q* directions to equal *σ*^2^. The log likelihood is given by:

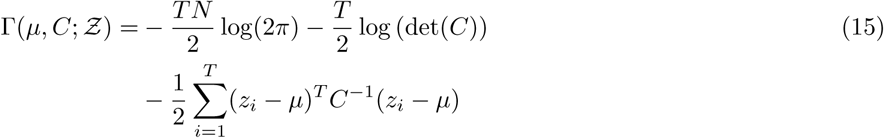

The maximum likelihood PPCA parameters have closed-form expressions that can be computed from an eigendecomposition of the sample covariance matrix. The mean is equal to the sample mean of *Ƶ*:

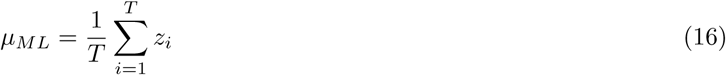

Let {λ_1_,…, λ_*N*_} denote the eigenvalues of the sample covariance matrix of *Ƶ* (in descending order) with corresponding eigenvectors {*u*_1_,…,*u*_*N*_}. The maximum likelihood noise variance is equal to the average of the smallest *N* − *q* eigenvalues:

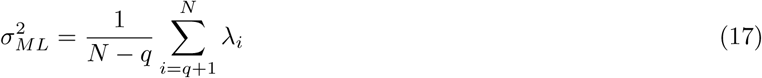

Let Λ_*q*_ = diag(λ_1_,…,λ_*q*_) and *U*_*q*_ = [*u*_1_,…, *u*_*q*_] contain the top *q* eigenvalues and eigenvectors of the sample covariance matrix. The maximum likelihood covariance matrix is:

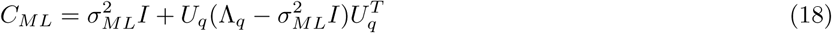

The dimensionality *q* is a hyperparameter, whose value can be selected based on the data. A separate value is selected for each local set of points on which PPCA is run. In our analyses, we chose *q* as the minimum number of dimensions containing fraction *α* = 0.95 of the total variance:

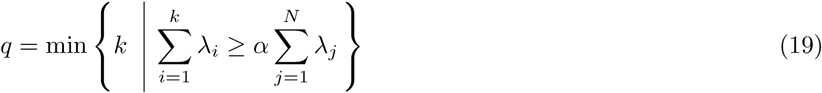

*q* can also be chosen using Bayesian model selection. In this case, *q* is selected to maximize the evidence *p*(*Ƶ* | *q*), for which^14^ gives an efficiently computable, closed form approximation.

### 2 Data processing

Analyses were performed on a subset of the data collected in.^15^ For each task (RFT, SMT), we selected 3 behavioral sessions to analyze, from 3 different rats. Individual sessions were chosen to maximize the number of simultaneously recorded cells, after those with very high or low firing rates were removed, as described in the following paragraph (cell count after removal, RFT sessions 52, 20, 41 for rats 103, 107, 108; SMT sessions 42, 15, 23 for rats 103, 106, 107).

After obtaining spike trains processed as in,^15^ we performed the following additional procedures. For each SMT session, we removed all spikes which occurred during the inter-trial interval (as defined in^15^), restricting our analyses to activity from the trial portion of the task. We then defined population activity vectors for each behavioral session via the following steps. We first removed cells with mean firing rate less than .05 Hz or greater than 5 Hz. The purpose of the first threshold was to reject SMT cells that had no activity during the trial-portion of task (which were the only cells removed), for the simplicity of implementing our analyses (including these cells would not have affected the results of the paper). The purpose of the second threshold was to remove putative interneurons.^15^ For each ofthe remaining cells, we computed a binned firing rate by convolving its spike train with a Gaussian kernel (standard deviation, .7 seconds), resulting in a smoothed, continuous-time signal which we sampled at a constant rate (temporal bin width, .05 seconds). We concatenated the firing rates in each time bin across cells to create population activity vectors, which were used in all subsequent analyses.

### 3 Reconstructed activity vectors and reconstruction score

In Sections 1.5.1 and 1.5.2 we defined a forward mapping *g*_*f*_ : ℝ^*N*^ → ℝ^*d*^ from state space to intrinsic manifold coordinates, and a reverse mapping *g*_*r*_ : ℝ^*d*^ → ℝ^*N*^ from intrinsic manifold coordinates to state space. The composition of these two maps defines a “reconstruction map” *h* = *g*_*r*_ º *g*_*f*_ : ℝ^*N*^ → ℝ^*N*^. For activity vector *x* ∈ ℝ^*N*^, let 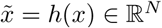 denote the corresponding reconstructed activity vector. The vector 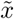 represents the image of *x* under a mapping from the activity vectors to the reconstructed manifold, viewed as a subspace of activity space.

The reconstruction score s (shown in Fig. 2b, and Extended Data Figs. 1 and 2) is defined as the fraction of variance in the activity vectors {*x*_*t*_} explained by the reconstructed activity vectors 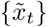. Specifically, if we define

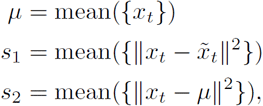

where ‖*x*‖ denotes the Euclidean norm of *x*, then s is given by

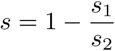

To compute the activity reconstructions shown in Fig. 3, we used a cross-validated version of the above procedure. For each trial *l*, we ran the MIND algorithm on a training set, defined as the union of activity vectors not from trial l, obtaining a forward map 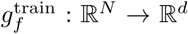 and a reconstruction map *h*^train^ : ℝ^*N*^ → ℝ^*N*^. We applied 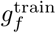 to the activity vectors from trial *l* (the test set) to produce a low-dimensional embedding of the network trajectory on that trial (as shown in Figs. 3d, 3e, top), and we applied *h*^train^ to produce a corresponding high-dimensional reconstructed trajectory (as shown in Figs. 3d, 3e, bottom, for two individual cells (corresponding to two coordinates of the high-dimensional reconstruction)). The scores associated with these reconstructions, as shown in Fig. 3c, are defined in Section 13.

### 4 Manifold firing fields

We use the terminology “manifold firing field” to refer to the activity of a given cell, viewed as a function of intrinsic manifold coordinates. Manifold firing fields were depicted in one of two ways.

Let *y* ∈ ℝ^*d*^ denote the intrinsic manifold coordinates of a particular high-dimensional state *x* ∈ ℝ^*N*^, and let *x*^*i*^ ∈ ℝ denote the value of the *i*th coordinate of *x*, i.e., the activity of cell *i* in that state. In Fig. 2e (SMT) as well as Extended Data Figs. 6 and 7, we depict manifold firing fields simply by visualizing *x*^*i*^ as a function of y. For aid of visualization, only landmark points are shown (see Section 1.6.3). In Fig. 2e (RFT), *x*^*i*^ is visualized as a function of the the first two manifold coordinates (ordered by variance). In Extended Data Figs. 6 and 7, all coordinates are shown.

In Figs. 3d and 3e, manifold firing fields are meant to indicate the activity reconstructed from the manifold. Therefore, these fields are not given by each cell’s true activity values *x*^*i*^, but rather its reconstructed activity values 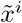 (see Section 3). Intuitively, these “reconstructed” manifold firing fields represent the activity of each cell projected onto, or smoothed along, the manifold. When the reconstruction is accurate, these fields will closely approximate the fields computed by the first procedure above.

### 5 Poisson model and behavioral firing fields

We tested the hypothesis that the hippocampal activity manifolds contained structure beyond the measured behavioral variables by comparing the real data to a Poisson null model.^16^ Under this model, each neuron’s activity was generated according to its behavioral firing field, and did not contain any sources of variability beyond Poisson noise.

To define behavioral firing fields for each cell, we divided the measured behavioral variables (position in the RFT or sound frequency in the SMT) into discrete bins, using bin sizes of 2 cm^2^ for position and .06 log kHz for frequency. The unsmoothed behavioral firing field 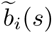 was given by the average firing rate of cell *i* in bin *s*. We smoothed 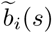 using a Gaussian kernel with standard deviation 1.5 times the bin size to produce the behavioral firing field *b*_*i*_(*s*).

Given the time series *s*(*t*) of behavioral variables measured in each session, we linearly interpolated the behavioral firing field *b*_*i*_(*s*) to define the time-varying rate *b*_*i*_(*s*(*t*)), which we denote *b*_*i*_(*t*) by abuse of notation. The synthetic Poisson spike train for cell *i* was then defined as a sample from an inhomogeneous Poisson process with underlying rate *b*_*i*_(*t*). We repeated this procedure for all cells, producing a simulated dataset which we analyzed exactly as the real data. For the dimensionality analysis in Fig. 2b, we used 100 simulated datasets.

### 6 Intrinsic dimensionality score

To estimate an optimal manifold dimensionality, we defined an intrinsic dimensionality score, adapted from the Gromov-Hausdorff distance between metric spaces.^17^ For each dimensionality *d*, the score *G* = *G*(*d*) was computed using the original data points 𝒳 = {*x*_*t*_} ⊂ ℝ^*N*^, their low-dimensional manifold coordinates 𝒴 = {*y*_*t*_} ⊂ ℝ^*N*^, and their reconstructions 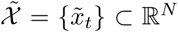. The score implemented a tradeoff between two terms, one quantifying how similar 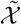 was to *𝒳*, the other quantifying how similar 𝒴 was to a uniform *d*-dimensional ball. In the expressions below, ‖*a*‖ denotes the Euclidean norm of vector *a*.

We first transformed the vectors in 𝒴 so they were on the same scale as the vectors in 𝒳, as follows. For a set of vectors *S* = {*s*_*t*_}, let denote the mean of *S*, and let *r*_*S*_ denote its radius, which we defined as the 95th percentile of the distances ‖*s*_*t*_ − *μ*_*S*_ ‖}. We scaled each element *y* ∈ 𝒴 by the transformation

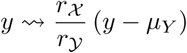

The score *G* was then defined as the negative of the maximum of two terms, *G* = −max(*G*_1_, *G*_2_) (we took the negative so that the optimal dimensionality occured as a peak rather than a trough, for the purpose of visualization).

We defined *G*_1_ to be the 95th percentile of the reconstruction errors 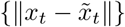.

To define *G*_2_, let *v*_*j*_ denote the *j*th coordinate vector in ℝ^*d*^ (i.e. the vector with *i*th coordinate equal to 1 if *i* = *j*, and 0 otherwise). For a vector *v*, let

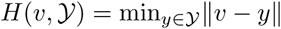

denote the minimum distance from v to a vector in 𝒴. We defined *G*_2_ to be the 95th percentile of the set of distances {*H*(±*v*_*j*_, 𝒴)} across all *v*_*j*_, 1 ≤ j ≤ *d*.

As *d* increases, *G*_1_ goes down as the reconstruction has more degrees of freedom for fitting the original data, but *G*_2_ goes up as the manifold coordinates of the activity vectors are less able to fill up space in the d-dimensional ball. We define the optimal manifold dimensionality to be the dimensionality which maximizes *G*(*d*).

Although the score was motivated by the Hausdorff and Gromov-Hausdorff distances between metric spaces,^17^ it does not involve any search over mappings between spaces; rather, we use the candidate mappings provided by MIND.

To display *G*(*d*) in Fig. 2b, we scale the scores for each dataset to range from 0 to 1, since we are only interested in the value of *G*(*d*) relative to the maximum.

In our tests, many details of the definition of *G*(*d*) could be changed, including changing the percentiles, or replacing them by the mean or median, without changing the optimal dimensionality estimates.

### 7 Peer-prediction score

As a cross-validated measure of reconstruction performance for each manifold dimensionality, we computed peer-prediction (or “leave-neuron-out”) scores as described in.^18,19^

The peer-prediction score quantifies the extent to which a given model can be used to predict the activity of one cell from the activity of its “peers” (the other recorded cells). For each dataset of activity vectors 𝒳, we computed the score as follows. The procedure requires cross-validation with respect to time as well as neurons. Therefore, we began by constructing k splits of the time points into training and test set, using the procedure described in Section 10. We took k = 4 for each dataset. We used these temporal splits for the rest of the analysis.

The analysis produces a single score *L*_*d*_ for each manifold dimensionality *d*, which itself is an average over scores, one for each choice of hold-out cell. For each choice of cell i, we removed cell i from the dataset, resulting in a set of *N* − 1-dimensional “peer” activity vectors 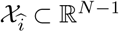, as well as the set 𝒳_*i*_ ⊂ ℝ of activity values for cell *i*. 𝒳_*i*_ can be thought of as a set of activity vectors for a population of size one.

For each choice of one of the *k* temporal splits, we applied the MIND algorithm to the training set 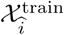 to produce intrinsic coordinates 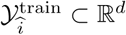, as well as a forward mapping 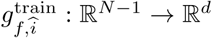. Then, using the same regression algorithm as for the forward and reverse mappings, we learned a “prediction mapping” 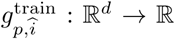, trained on input-output pairs from 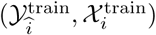. The prediction mapping reconstructs the activity of the held-out cell from the activity of its peers. It was trained on landmark points only, as in the original algorithm.

Let *L* denote a choice of loss function, and let j be an index denoting the choice of temporal split. We computed the loss for each temporal-split:

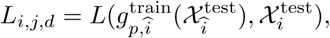

which we used to compute the test loss for cell *i* (and manifold dimensionality *d*):

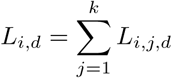

We took *L*(*x, y*) = ‖*x* − *y*‖^2^*/T* (where *x* and *y* are scalar time series of length *T*), and we then defined the peer-prediction score *L*_*d*_ to be the mean Euclidean error for each cell,

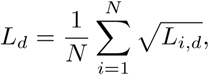

so that the final loss funtion was a mean root mean squared error (RMSE) across cells.

Finally, for computational reasons, the analysis was restricted to the 30 cells from each behavioral session with the highest maximum firing rate (if the session had fewer than 30 cells, all cells were used).

### 8 Field stability plots

To examine field stability we split the rat 103 SMT dataset into its first and second halves. We then used MIND to separately fit a 2d manifold to each half (2 being the optimal dimension, as previously estimated), and we assessed stability by plotting the firing fields from the first and second half side by side for each cell. Fields were plotted for the 15 cells with the highest firing rate across the session.

We marked the center of mass of each field in red. If 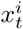 denotes the firing rate of the *i*th cell at time t, LM denotes the set of time points corresponding to landmark points, and *y*_*t*_ denotes the 2d manifold coordinates for the state at time *t*, then the center of mass *c*_*i*_ for the ith cell is

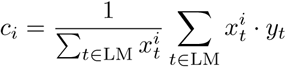

### 9 Simulated high-dimensional place cells

To validate our intrinsic dimensionality estimator, we constructed a set of simulated high-dimensional datasets, modeled as idealized place cells for a rat foraging in a high-dimensional arena. Any dimensionality estimator will be subject to the “curse of dimensionality,” in that the sample complexity of the dimensionality estimation problem will generally be exponential in the true dimensionality. Therefore, in addition to idealized, highly-sampled simulations to validate the definition and implementation of the estimator, we also tested the estimator on simulations with sampling matched to the data.

For each dimensionality *d*, a simulated dataset consisted of a set of firing rates over time, defined in terms of a trajectory {*x*_*t*_} in the *d*-dimensional unit ball *B* (i.e., the set of points in ℝ^*d*^ with norm less than or equal to 1). The firing rate of cell *i* was determined by its firing field, which was modeled as a *d*-dimensional Gaussian bump with center *c*_*i*_ ∈ ℝ^*d*^, amplitude *a*_*i*_ ∈ ℝ, and standard deviation *σ*_*i*_ ∈ ℝ. At time t, the firing rate of cell *i* was given by

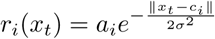

The random trajectory {*x*_*t*_} was generated as follows. We first chose a set of m unordered vectors {*x*} uniformly at random from B. We created a trajectory by ordering these vectors via an iterative procedure, defined in terms of a parameter s which specified the approximate step size. The procedure was initialized by choosing *x*_1_ at random from {*x*}. Next, given an ordering *x*_1_, …,*x*_*t*_ up to time *t*, we defined a target *s*_*t*_ for the *t*th step size (i.e., the distance ‖*x*_*t*_ − *x*_*t*+1_‖) by taking *s*_*t*_ = *ϵ*_*t*_*s*, where *ϵ*_*t*_ was chosen randomly from [0,1] at each step. The next vector *x*_*t*+1_ was then chosen to minimize the value

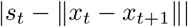

subject to the following criteria: 1) *x*_*t*+1_ was not already chosen in the sequence; 2) *x*_*t*+1_ was closer to *x*_*t*_ than to *x*_*t*-1_, i.e.,

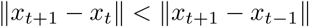

This last criterion encouraged the trajectory to move forwards; if no remaining vectors satisfied it, it was ignored. Finally, we linearly interpolated this sequence of *m* vectors to reach a target trajectory length of *T* time steps. We verified that the trajectories had no unrealistic discontinuities by ensuring that the largest step size was less than 2*s*.

In all simulations, we took *s* = .1, *m* = 5000, and *T* = 17500 (the number of time steps in the rat 103 RFT dataset).

For the highly-sampled idealized simulations, we used 128 cells, with *a*_*i*_ = 1 and *σ*_*i*_ = .5 for each cell. We defined centers *c*_*i*_ to be vertices of the unit cube centered at the origin, chosen randomly without replacement. When there were more than 2^*d*^ centers, we chose the remaining centers uniformly at random from *B*.

For the data-matched simulations, we matched the cell parameters to properties of the rat 103 RFT dataset. We used the same number of cells (52), and for the *i*th cell, we set *a*_*i*_ equal to the maximum firing rate of the *i*th cell in the dataset. We chose field centers *c*_*i*_ independently and uniformly at random from *B*. Next, having chosen *a*_*i*_ to match each cell’s maximum firing rate, we adjusted *σ*_*i*_ in order to match each cell’s mean. Specifically, after choosing *a*_*i*_, *c*_*i*_ and computing the trajectory as above, a choice of *σ*_*i*_ determined the firing rate *r*_*i*_(*x*_*t*_). We therefore used a 1d bounded optimization routine to find the value of *σ*_*i*_ in the interval [0, 2] which minimized the distance between mean(*r*_*i*_(*x*_*t*_)) and the mean of the *i*th cell’s true firing rate. We implemented this optimization with Matlab’s `fminbnd` function. Finally, we applied the same Poisson noise procedure as described in Section 5, generating smoothed Poisson spike trains from the underlying simulated firing rates.

For all simulations, we ran the MIND algorithm with the same parameters as for the data.

### 10 Decoding measured behavioral variables

We performed decoding using cross-validated Gaussian process regression (GPR). Our inputs were a set of vectors {*r*(*t*)} over time, either low-dimensional embedding coordinates 𝒴, or network states 𝒳. The outputs to be decoded were behavioral variables {*s*(*t*)}.

To minimize the influence of local temporal correlations, we employed a blocked cross-validation approach. We first divided time into k contiguous blocks of equal size, and we removed the last *T*_buff_ = 1 second from each block. For the *i*th cross-validation fold, *i* = 1,…,*k*, the *i*th block was held out as the test set, and the remaining blocks were combined to form the training set. We then trained a GPR model to predict *s*(*t*) from *r*(*t*) using landmark points from the training set (we trained on landmark points for computational reasons; see Section 1.6.3 for their definition). In the RFT, where *s*(*t*) was the animal’s position, we treated *x* and *y* positions separately, fitting independent models to predict each coordinate. The model fits were performed using the `fitrgp` function in Matlab 2018a, with default parameters (i.e., using the squared exponential kernel, with hyperparameters fit by maximizing the marginal likelihood). For each fold, we applied the trained model to the test set, and defined the test set error to be the average Euclidean distance between the true and predicted behavioral variables. We defined the final decoding error to be the average of the test errors across all *k* folds.

The error bars in Fig. 2c indicate the standard deviation of the *k* test set errors. We took *k* = 10 in our analyses.

### 11 Analyzing single-trial activity in the SMT

In the SMT, we analyzed single-trial activity by selecting peaks in the firing rate on each trial, using the following procedure.

We analyzed each SMT dataset separately. We first removed all cells that fired fewer than 5 spikes per trial, unless this left less than 10 cells, in which case we took the 10 cells with the highest maximum firing rate (for rats 103, 106, and 107, this resulted in 11, 10, and 10 cells, respectively).

For a fixed cell and trial, we defined a set of significant peaks on that trial. Significant peaks were defined as peaks in the firing rate vs. log frequency curve *r* = *r*(*f*) satisfying two criteria: 1) they exceeded mean(*r*) + s.d.(*r*), and 2) they had a peak prominence exceeding mean(*r*)/2, meaning there was a vertical drop of at least mean(*r*)/2 on each side of the peak before another peak or the edge of the signal was reached. We implemented this using the `findpeaks` function in Matlab 2018a, after zero padding the signal *r* on the left and right to ensure that edge points could be included as peaks.

In what follows, as well as in the main text, we refer to the location (i.e, frequency value) of a peak as its *position,* and we refer to the height of a peak as its *amplitude.* For simplicity, we normalized the vector *f* of log frequency bins to the interval [0,1] by subtracting its minimum value and then dividing by its maximum. Furthermore, for each cell, we scaled *r*(*f*) by its maximum value across trials. Therefore, peak positions and amplitudes each ranged between 0 and 1.

For simplicity, we analyzed trial-to-trial variability with respect to one primary peak location, and did not consider multiple peaks. Therefore, for each cell and trial, we needed to choose a primary peak from the set of significant peaks on that trial. However, since our goal was to quantify peak variability, we needed to avoid artifactual variability which would arise from cells with multiple peaks, if we selected mismatched primary peaks from trial to trial. To this end, for each cell, we simply chose the assignment of primary peaks to trials which made position variability as small as possible. Algorithmically, we implemented this as follows: for a given frequency bin *f*, we took the primary peaks on each trial to be the significant peaks with position closest to *f* (including no selection, if the trial had no significant peaks). For each *f*, we combined these primary peaks into a set *S*, and computed its position variability score, defined as the interdecile range of the positions of all peaks in *S*. We iterated this procedure over all frequency bins *f*, and took the set S with the smallest variability score as the set of trial-by-trial peaks for cell *i*.

### 12 Quantification of trial-to-trial variability

The procedure in Section 11 defined a set *S* of primary peaks for each cell *i*, with at most one per trial. We refer to the sets of positions and amplitudes of the peaks in *S* as the position and amplitude distributions for cell *i*, respectively, as shown in Fig. 3b (top). We quantified variability for each cell by computing the interdecile range of its position and amplitude distributions, as shown in Fig. 3b (bottom).

### 13 Single-trial reconstruction scores

To quantify how well the peak positions and amplitudes from the reconstructed activity matched those in the data, we first selected peaks from the reconstructed activity 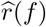 as follows (see Section 3 for the reconstruction procedure itself).

We selected significant peaks on each trial, as in Section 11. For a given trial *j*, let *Ŝ*_*j*_ denote the significant peaks for the reconstructed firing rate on trial *j*, and let *S*_*j*_ denote the significant peaks for the real firing rate. Let *s*_*j*_ ∈ *S*_*j*_ denote the primary peak, as described in Section 11. To pair *s*_*j*_ with a corresponding reconstructed primary peak *ŝ*_*j*_ ∈ *Ŝ*_*j*_, we used the Hungarian algorithm^20^ to match elements of *S*_*j*_ and *Ŝ*_*j*_. We then defined *ŝ*_*j*_ to be the element assigned to *s*_*j*_, or to be empty (i.e., no reconstructed peaks on that trial) if no element was assigned to *s*_*j*_. We used this matching approach as a way of handling the existence of multiple peaks in the activity and its reconstruction, given our goal of analyzing a single peak per trial. Rather than simply pairing *s*_*j*_ with its nearest reconstructed peak, this more conservative procedure paired *s*_*j*_ with the nearest reconstructed peak that was not more preferably matched to another significant peak in the real activity. However, follow-up analyses might treat multiple peaks more directly.

For each cell, once we had a choice of reconstructed primary peak *ŝ*_*j*_ for every trial, we quantified the agreement between the real and reconstructed peaks by computing the mean absolute error (MAE) between real and reconstructed positions and amplitudes across all *R* trials. For real and reconstructed positions 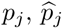, this is

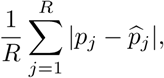

and likewise for amplitudes. On less than 3% of trials, either the real or reconstructed firing rate had no primary peak, in which case that trial was removed and did not contribute to the error.

### 14 Noise correlations

We computed noise correlations as in^21,22^. For cell i recorded during the SMT, let *b*_*i*_(*s*) denote its behavioral firing field (see Section 5), which gives the expected activity of cell *i* at sound frequency *s*. If *s*(*t*) denotes the time course of frequency throughout the behavioral session, then we let *b*_*i*_(*t*) = *b*_*i*_(*s*(*t*)) denote the expected activity given the frequency value at time *t*, by abuse of notation. If *r*_*i*_(*t*) denotes the firing rate of cell *i* as a function of time, we define the noise correlation *c*_*ij*_ as the Pearson correlation coefficient between 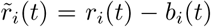 and 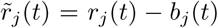, thought of as vectors indexed by time. In notation, this is

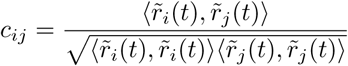

where ⟨ *x, y* ⟩ denotes the inner product of vectors *x* and *y*. As a summary statistic in Fig. 3f, we compute the average absolute value of the noise correlations across all cell pairs.

### 15 Noise correlation shuffles

To test for noise correlations in the SMT, we defined a shuffle procedure that preserved the average sound frequency tuning of every cell, while removing any trial-by-trial correlations between cells beyond those resulting from their frequency tuning.

Conceptually, the shuffle is implemented by replacing a cell’s activity on trial *a* by its activity on trial *b*, chosen at random, after warping time on trial b so that the frequency sweeps in both trials are aligned. This is repeated for each trial, and the same shuffle is then repeated independently for each cell.

In detail, let 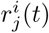 denote the firing rate of cell *i* on trial *j* as a function of time *t*, and let *f*_*j*_(*t*) denote sound frequency as a function of time on the same trial. Choose a random permutation of trial labels, *j* ↦ *j*′. We will replace the activity vector 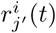 on trial *j*′ by the activity vector 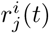 on trial *j*, while preserving the frequency dependency of 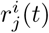. To do so, we consider 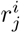 as a function of *f* = *f*_*j*_(*t*) (*f*_*j*_(*t*) and *f*_*j*′_(t) are strictly increasing frequency sweeps through the same frequency values). We linearly interpolate 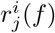 at query values *f*_*j*′_(t) to create a new activity vector 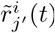. We do this for all trials, and concatenate the 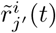 to create a shuffled activity vector 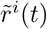 for cell *i*. To create a shuffled dataset, we repeated this procedure independently for all cells. We used 250 noise correlation shuffles in our analyses.

We remark that this shuffle procedure is feasible due to the trial structure of the SMT, in which the same sequence of measured behavioral variables is presented on every trial, up to temporal rescaling. An analogous shuffle in the RFT, for example, would be less straightforward.

### 16 Synthetic place and grid cell data

Synthetic place and grid cell activity was used to validate manifold discovery methods. This data was generated using a model network containing 50 place cells and 50 grid cells, driven by the simulated trajectory of a rat exploring a two-dimensional box with side length 88 cm. The instantaneous firing rate of each cell was given by the amplitude of its place or grid field at the current position of the rat, with additive white Gaussian noise (noise standard deviation was 1/5th the signal standard deviation).

Place fields were modeled as isotropic Gaussian functions centered randomly in the environment, with width 41.6 cm (full width at half maximum; FWHM). Each grid field was modeled as a sum of Gaussian bumps centered on a hexagonal lattice. Grid cells were organized in four modules, and grid fields within a module shared a characteristic spacing, width, and angle. The lattice spacing and bump width for each module increased by a factor of 1.3 relative to the previous module, with base spacing 39.8 cm and base width 27.4 cm (FWHM). The angle of the hexagonal lattice progressed in increments of 15° across modules. Lattice phases were random for each grid cell. These parameters were chosen to approximately match previous experimental observations.^23^

### 17 Deviation analysis

#### 17.1 Predicting behavior from manifold coordinates

To identify deviations from coding instantaneous behavior in the RFT, we predicted the rat’s position from the 3d manifold coordinates representing population activity at each timepoint. Time series containing position and manifold coordinates were first downsampled to a rate of 4 Hz to speed up subsequent Monte Carlo analyses. Cross-validation was used to avoid overfitting during position decoding, and was performed in a blocked manner to minimize temporal dependence between training and held-out data. The time series containing manifold coordinates and animal position were partitioned into ten contiguous blocks, and each block was held out in turn. A Gaussian process regression model was fit to the remaining data, then used to predict position from manifold coordinates in the held out block (see Section 10 for details). Predictions for all held-out blocks were then concatenated to give a time series 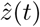, specifying the predicted position at time *t*.

#### 17.2 Comparison to Poisson model

When analyzing deviations, we used the Poisson model as a null model. In this case, activity encodes the rat’s instantaneous position by construction, and any difference between the predicted and actual trajectories through the environment is a consequence of random variation. We fit the Poisson model and generated synthetic activity as in Section 5. 2000 surrogate datasets were generated for each rat, containing firing rates along the rat’s actual trajectory through the environment. A manifold (with dimensionality 3, chosen to match the real data) was fit to the expected firing rates (i.e. the rate function of the Poisson process for each cell) as in Section 1. The forward mapping for this manifold was used to obtain 3d manifold coordinates for each surrogate dataset, as in Section 1.5.1. Manifold coordinates were used to predict position as described above for the real data, using two-fold blocked cross-validation.

#### 17.3 Lag estimation

Leading and lagging deviations were quantified by estimating the temporal lag between the predicted position z(t) and actual position 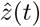 at each timepoint, using dynamic time warping (DTW).^24^ DTW aligns two time series by nonuniformly stretching and shrinking them along the time axis to minimize a distance measure. In our analyses, DTW was used to align *z* and 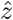, producing a warping path *W* = {(*u*_*i*_, *v*_*i*_)} that minimized the Euclidean distance between points in the warped time series:

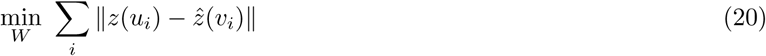

The warping path *W* consists of a series of paired time points such that *z*(*u*_*i*_) is matched to 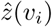, with time points constrained to increase continuously and monotonically, and to cover the entire range of the time series. The warping path was also constrained by requiring that |*u*_*i*_ −*v*_*i*_| did not exceed 5 s.

Lag was estimated as the time elapsed between the actual position at time t and the predicted position assigned to it by DTW (or the mean time elapsed if there were multiple assignments). Positive lag indicated that *z* was leading *z*, and negative lag indicated that it was lagging behind.

For all subsequent analyses involving estimated lags, we included only timepoints where the actual position matched its DTW-assigned predicted position with high confidence. To meet this criterion, the actual position must fall within the 75% credible region for the corresponding predicted position—the central, ellipsoidal region of the Gaussian posterior distribution (given by Gaussian process regression) containing 75% of the probability mass. This criterion excluded points where the trajectories did not reliably match, such as detour deviations.

#### 17.4 Lag distribution

Fig. 4b and Extended Data Fig. 2 show the overall distribution of estimated lags across time. The probability density function was estimated using a kernel density estimator (Gaussian kernel, bandwidth selected using two-fold blocked cross-validation). The cumulative distribution function was estimated as the empirical CDF.

The overall magnitude of estimated lead and lag was quantified as the mean absolute lag across timepoints. For the Poisson model, we computed the mean and standard error of this statistic across surrogate datasets. To test the hypothesis that mean absolute lag was greater for the real data than the Poisson model, we calculated a one-tailed p value as the fraction of surrogate datasets where mean absolute lag was greater than or equal to that in the real data.

#### 17.5 Significant leads & lags

We sought to identify individual timepoints where lead or lag was unlikely to have arisen due to random variation as in the Poisson model. At each timepoint, we tested the hypothesis that the real data had lag of greater magnitude than the Poisson model. Two-tailed p values were calculated as the fraction of surrogate datasets where the absolute lag was greater than or equal to that in the real data. To account for multiple comparisons over many timepoints, we controlled the false discovery rate (FDR) at a level of 0.05, using the procedure described in.^25^ Significant leads and lags were identified as timepoints with positive or negative lag, respectively, and FDR q value less than 0.05. Because the rat’s trajectory was identical for the real data and Poisson model, comparing individual timepoints controlled for any influence of local trajectory properties on the estimated lag.

#### 17.6 Detour detection

Detour deviations occurred when the actual position differed strongly from the predicted position, and this difference could not be accounted for by lead or lag. To detect detours, we compared the actual position at each timepoint to its DTW-assigned predicted position. The use of DTW assignments excluded differences due to lead or lag. Detours were said to occur when the actual position fell outside the 95% credible region for the corresponding predicted position— the central, ellipsoidal region of the Gaussian posterior distribution (given by Gaussian process regression) containing 95% of the probability mass.

#### 17.7 Analysis of manifold reconstruction error

To determine whether population activity remains on the manifold during deviations, we examined the manifold reconstruction error—the Euclidean distance in firing rate space between population activity and its projection on the manifold (i.e, its image under the reconstruction map, see Section 3). We calculated the Pearson correlation of reconstruction error with decoding error, and with absolute lag. 95% confidence intervals were estimated using a moving block bootstrap approach (10000 iterations) to account for temporal dependence.^26,27^ Stationarity of the time series to be bootstrapped was first verified using augmented Dickey-Fuller and Phillips-Perron tests with multiple lags and significance level *α* = 0.05. Block size for the moving block bootstrap should be large enough to cover the range of temporal dependence in the data. It was chosen to be at least twice the lag at which the autocorrelation function of any time series decayed to a level that was statistically indistinguishable from zero (with *α* = 0.05). Changing the block size did not affect our results (tested within a factor of 2).

### 18 Comparison to PCA and isomap

In Extended Data Fig. 5 we compared MIND to principal component analysis (PCA) and isomap,^7^ two well-known methods for manifold learning and dimensionality reduction. We chose these methods because of their widespread use, and because MIND builds directly on ideas from both techniques. In addition to other manifold learning algorithms,^6,18,28-37^ all three methods serve complementary purposes, and are useful in different situations.

A fundamental similarity between MIND, PCA, and isomap is that they produce a low-dimensional representation of the data by attempting to preserve distances.^6^ All three methods effectively use a form of multidimensional scaling (MDS)^10^ to obtain low-dimensional manifold coordinates where Euclidean distances match distances computed between the original data points. PCA and isomap use classical MDS, whereas MIND uses Sammon mapping^9^-a variant of nonclassical, metric MDS that places greater emphasis on preserving small distances.

However, each technique measures distances between data points in a unique way. PCA uses Euclidean distances, whereas isomap uses geodesic distances, which are estimated from local Euclidean distances between neighboring points. MIND uses a novel distance measure based on estimated transition probabilities between network states (see section 1.3). For each method, the distance measure is an essential choice governing the type of structure to capture.

For example, as a consequence of their distance measures, isomap and MIND can discover nonlinear manifolds, whereas PCA is restricted to the linear case. However, PCA is more efficient when the underlying structure is truly linear. Another important distinction is that MIND captures structure in the dynamics of the system, whereas PCA and isomap are insensitive to dynamics-randomly permuting the temporal order of the data would not change their output. In contrast, PCA and isomap are sensitive to the relative positions of data points in input space, as a consequence of their use of Euclidean distances. For example, changing the firing rate of a neuron might strongly affect the distances computed between network states. However, the MIND distance measure does not directly depend on the firing rates in different states, but rather how the system moves through these states. In this manner, MIND attempts to learn natural distances from the behavior of the network itself, rather than ascribe *a priori* meaning to particular differences in activity patterns.

To compare methods quantitatively, we fit 2d manifolds to population activity recorded from rat 103 in the RFT. Because isomap does not provide a mapping from intrinsic manifold coordinates back to activity space, we asked how well each manifold captured the measured behavioral variables. Given that position in the 2d environment accounts for a large portion of the variation in firing rates over time, a 2d manifold that captures this structure should be similar to the environment. To visualize results, we plotted manifold coordinates colored by the rat’s position in the environment at each timepoint (Extended Data Fig. 5a). The manifold recovered by MIND smoothly encoded position, whereas dissimilar positions were more intermixed on manifolds recovered by isomap and PCA. To quantify the geometric similarity between manifold and environment, we compared the Euclidean distance between every pair of points on the manifold to the distance between the corresponding pair of points in the environment (Extended Data Fig. 5b). Pearson correlation between distances was 0.76, 0.55, and 0.23 for MIND, isomap, and PCA, respectively (2 × 10^6^ pairs of points). Finally, to measure how well the manifold captured information about position, we decoded position from manifold coordinates as in section 10. Extended Data Fig. 5c shows mean squared decoding error, estimated using 10 fold cross-validation over temporally contiguous blocks of data. Decoding error was 1.8%, 57.7%, and 116.6% greater than the raw data for MIND, isomap, and PCA, respectively.

Therefore, with dimensionality matched to the number of measured behavioral variables, the manifold recovered by MIND is geometrically similar to the environment. Furthermore, it captures most of the information about the rat’s position that is available to the decoder in the raw firing rates. In contrast, isomap and PCA recover manifolds that are less similar to the environment, and capture less information about position. Determining how such performance differences relate to the various conceptual differences between methods is an interesting topic for future investigation.

## References

1. Moser, M.-B., Rowland, D. C. & Moser, E. I. Place cells, grid cells, and memory. Cold Spring Harbor Perspectives in Biology 7 (2015).

2. Pastalkova, E., Itskov, V., Amarasingham, A. & Buzsaki, G. Internally generated cell assembly sequences in the rat hippocampus. Science 321, 1322–1327 (2008).

3. Aronov, D., Nevers, R. & Tank, D. W. Mapping of a non-spatial dimension by the hippocampal-entorhinal circuit. Nature 543, 719–722 (2017).

4. O’Keefe, J. & Nadel, L. The hippocampus as a cognitive map (Oxford: Clarendon Press, 1978).

5. Schiller, D. et al. Memory and space: Towards an understanding of the cognitive map. Journal of Neuroscience 35, 13904–13911 (2015).

6. Fenton, A. A. & Muller, R. U. Place cell discharge is extremely variable during individual passes of the rat through the firing field. Proceedings of the National Academy of Sciences 95, 3182–3187 (1998).

7. Cohen, M. R. & Kohn, A. Measuring and interpreting neuronal correlations. Nature Neuroscience 14, 811–819 (2011).

8. Arieli, A., Sterkin, A., Grinvald, A. & Aertsen, A. Dynamics of ongoing activity: explanation of the large variability in evoked cortical responses. Science 273, 1868–1871 (1996).

9. Renart, A. & Machens, C. K. Variability in neural activity and behavior. Current Opinion in Neurobiology 25, 211–220 (2014).

10. Doiron, B., Litwin-Kumar, A., Rosenbaum, R., Ocker, G. K. & Josic, K. The mechanics of state-dependent neural correlations. Nature Neuroscience 19, 383–393 (2016).

11. Johnson, A., Fenton, A. A., Kentros, C. & Redish, A. D. Looking for cognition in the structure within the noise. Trends in Cognitive Sciences 13, 55–64 (2009).

12. Redish, A. D. Vicarious trial and error. Nature Reviews Neuroscience 17, 147–159 (2016).

13. Harris, K. D., Csicsvari, J., Hirase, H., Dragoi, G. & Buzsaki, G. Organization of cell assemblies in the hippocampus. Nature 424, 552–556 (2003).

14. Carr, M. F., Jadhav, S. P. & Frank, L. M. Hippocampal replay in the awake state: a potential substrate for memory consolidation and retrieval. Nature Neuroscience 14, 147–153 (2011).

15. Hopfield, J. J. Neurodynamics of mental exploration. Proceedings of the National Academy of Sciences 107, 1648–1653 (2010).

16. Pfeiffer, B. E. & Foster, D. J. Hippocampal place-cell sequences depict future paths to remembered goals. Nature 497, 74–79 (2013).

17. Dayan, P. Improving generalization for temporal difference learning: the successor representation. Neural Computation 5, 613–624 (1993).

18. Stachenfeld, K. L., Botvinick, M. M. & Gershman, S. J. The hippocampus as a predictive map. Nature Neuroscience 20, 1643–1653 (2017).

19. McNaughton, B. L. et al. Deciphering the hippocampal polyglot: the hippocampus as a path integration system. Journal of Experimental Biology 199, 173–185 (1996).

20. Wimmer, K., Nykamp, D. Q., Constantinidis, C. & Compte, A. Bump attractor dynamics in prefrontal cortex explains behavioral precision in spatial working memory. Nature Neuroscience 17, 431–439 (2014).

21. Seung, H. S. How the brain keeps the eyes still. Proceedings of the National Academy of Sciences 93, 13339–13344 (1996).

22. Zhang, K. Representation of spatial orientation by the intrinsic dynamics of the head-direction cell ensemble: a theory. Journal of Neuroscience 16, 2112–2126 (1996).

23. Mazor, O. & Laurent, G. Transient dynamics versus fixed points in odor representations by locust antennal lobe projection neurons. Neuron 48, 661–673 (2005).

24. Galan, R. F., Sachse, S., Galizia, C. G. & Herz, A. V. M. Odor-driven attractor dynamics in the antennal lobe allow for simple and rapid olfactory pattern classification. Neural Computation 16, 999–1012 (2004).

25. Gallego, J. A., Perich, M. G., Miller, L. E. & Solla, S. A. Neural manifolds for the control of movement. Neuron 94, 978–984 (2017).

26. van der Maaten, L., Postma, E. O. & van den Herik, H. J. Dimensionality reduction: A comparative review. Tech. Rep. TiCC-TR 2009-005, Tilburg University (2009).

27. Yu, B. M. et al. Gaussian-process factor analysis for low-dimensional single-trial analysis of neural population activity. Journal of Neurophysiology 102, 614–635 (2009).

28. Churchland, M. M. et al. Neural population dynamics during reaching. Nature 487, 51–56 (2012).

29. Chen, Z., Gomperts, S. N., Yamamoto, J. & Wilson, M. A. Neural representation of spatial topology in the rodent hippocampus. Neural Computation 26, 1–39 (2013).

30. Cunningham, J. P. & Yu, B. M. Dimensionality reduction for large-scale neural recordings. Nature Neuroscience 17, 1500–1509 (2014).

31. Park, M. et al. Bayesian manifold learning: the locally linear latent variable model (ll-lvm). In Advances in Neural Information Processing Systems, 154–162 (2015).

32. Gao, Y., Archer, E. W., Paninski, L. & Cunningham, J. P. Linear dynamical neural population models through nonlinear embeddings. Advances in Neural Information Processing Systems 29, 163–171 (2016).

33. Linderman, S. W., Johnson, M. J., Wilson, M. A. & Chen, Z. A Bayesian nonparametric approach for uncovering rat hippocampal population codes during spatial navigation. Journal of Neuroscience Methods 263, 36–47 (2016).

34. Sussillo, D., Jozefowicz, R., Abbott, L. & Pandarinath, C. LFADS - latent factor analysis via dynamical systems. arXiv (2016). https://arxiv.org/abs/1608.06315.

35. Wu, A., Roy, N. G., Keeley, S. & Pillow, J. W. Gaussian process based nonlinear latent structure discovery in multivariate spike train data. Advances in Neural Information Processing Systems 30, 3496–3505 (2017).

36. Fox, S. E. & Ranck Jr, J. B. Improved method for measuring the topological dimensions of neuronal firing rate space. Program No. 616.12. In 2017 Neuroscience Meeting Planner (Washington, DC: Society for Neuroscience, 2017). Online.

37. Wei, Z., Inagaki, H., Li, N., Svoboda, K. & Druckmann, S. An orderly single-trial organization of population dynamics in premotor cortex predicts behavioral variability. bioRxiv (2018). https://www.biorxiv.org/content/ early/2018/07/25/376830.

38. Williams, A. H. et al. Unsupervised discovery of demixed, low-dimensional neural dynamics across multiple timescales through tensor component analysis. Neuron 98, 1099–1115.e8 (2018).

39. Muller, R. U. & Kubie, J. L. The firing of hippocampal place cells predicts the future position of freely moving rats. Journal of Neuroscience 9, 4101–4110 (1989).

40. Dabaghian, Y., Brandt, V. L. & Frank, L. M. Reconceiving the hippocampal map as a topological template. eLife 3, e03476 (2014).

41. Brody, C. D. & Hanks, T. D. Neural underpinnings of the evidence accumulator. Current Opinion in Neurobiology 37, 149–157 (2016).

42. Ziv, Y. et al. Long-term dynamics of CA1 hippocampal place codes. Nature Neuroscience 16, 264 (2013).

## References

1. Tipping, M. E. & Bishop, C. M. Probabilistic principal component analysis. Journal of the Royal Statistical Society: Series B (Statistical Methodology) 61, 611–622 (1999).

2. Dasgupta, S. & Freund, Y. Random projection trees and low dimensional manifolds. In Proceedings of the Fortieth Annual ACM Symposium on Theory of Computing, 537–546 (ACM, 2008).

3. Kpotufe, S. & Dasgupta, S. A tree-based regressor that adapts to intrinsic dimension. Journal of Computer and System Sciences 78, 1496–1515 (2012).

4. Breiman, L. Random forests. Machine Learning 45, 5–32 (2001).

5. Friedman, J., Hastie, T. & Tibshirani, R. The elements of statistical learning, vol. 1 (Springer Series in Statistics, New York, 2001).

6. van der Maaten, L., Postma, E. O. & van den Herik, H. J. Dimensionality reduction: A comparative review. Tech. Rep. TiCC-TR 2009-005, Tilburg University (2009).

7. Tenenbaum, J. B., De Silva, V. & Langford, J. C. A global geometric framework for nonlinear dimensionality reduction. Science 290, 2319–2323 (2000).

8. Johnson, D. B. Efficient algorithms for shortest paths in sparse networks. Journal of the ACM 24, 1–13 (1977).

9. Sammon, J. W. A nonlinear mapping for data structure analysis. IEEE Transactions on Computers 100, 401–409 (1969).

10. Borg, I. & Groenen, P. J. Modern multidimensional scaling: Theory and applications (Springer Science & Business Media, 2005).

11. Saul, L. K. & Roweis, S. T. Think globally, fit locally: unsupervised learning of low dimensional manifolds. Journal of Machine Learning Research 4, 119–155 (2003).

12. Torgo, L. F. R. A. Inductive learning of tree-based regression models. Ph.D. thesis, Universidade do Porto, Reitoria (1999).

13. De Silva, V. & Tenenbaum, J. B. Sparse multidimensional scaling using landmark points. Tech. Rep., Stanford University (2004).

14. Minka, T. P. Automatic choice of dimensionality for pca. Advances in Neural Information Processing Systems 598–604 (2001).

15. Aronov, D., Nevers, R. & Tank, D. W. Mapping of a non-spatial dimension by the hippocampal-entorhinal circuit. Nature 543, 719–722 (2017).

16. Fenton, A. A. & Muller, R. U. Place cell discharge is extremely variable during individual passes of the rat through the firing field. Proceedings of the National Academy of Sciences 95, 3182–3187 (1998).

17. Gromov, M. Metric structures for Riemannian and non-Riemannian spaces, Chapter 3 (Springer Science & Business Media, 2007).

18. Yu, B. M. et al. Gaussian-process factor analysis for low-dimensional single-trial analysis of neural population activity. Journal of Neurophysiology 102, 614–635 (2009).

19. Harris, K. D., Csicsvari, J., Hirase, H., Dragoi, G. & Buzsaki, G. Organization of cell assemblies in the hippocampus. Nature 424, 552 (2003).

20. Munkres, J. Algorithms for the assignment and transportation problems. Journal of the Society for Industrial and Applied Mathematics 5, 32–38 (1957).

21. Cohen, M. R. & Kohn, A. Measuring and interpreting neuronal correlations. Nature Neuroscience 14, 811 (2011).

22. Peyrache, A., Lacroix, M. M., Petersen, P. C. & Buzsaki, G. Internally organized mechanisms of the head direction sense. Nature Neuroscience 18, 569 (2015).

23. Stensola, H. et al. The entorhinal grid map is discretized. Nature 492, 72 (2012).

24. Berndt, D. J. & Clifford, J. Using dynamic time warping to find patterns in time series. In KDD Workshop, vol. 10, 359–370 (Seattle, WA, 1994).

25. Storey, J. D. A direct approach to false discovery rates. Journal of the Royal Statistical Society: Series B (Statistical Methodology) 64, 479–498 (2002).

26. Davison, A. C. & Hinkley, D. V. Bootstrap methods and their application, vol. 1 (Cambridge University Press, 1997).

27. Lahiri, S. N. Resampling methods for dependent data (Springer Science & Business Media, 2013).

28. Gao, Y., Archer, E. W., Paninski, L. & Cunningham, J. P. Linear dynamical neural population models through nonlinear embeddings. Advances in Neural Information Processing Systems 29, 163–171 (2016).

29. Wu, A., Roy, N. G., Keeley, S. & Pillow, J. W. Gaussian process based nonlinear latent structure discovery in multivariate spike train data. Advances in Neural Information Processing Systems 30, 3496–3505 (2017).

30. Linderman, S. W., Johnson, M. J., Wilson, M. A. & Chen, Z. A Bayesian nonparametric approach for uncovering rat hippocampal population codes during spatial navigation. Journal of Neuroscience Methods 263, 36–47 (2016).

31. Chen, Z., Gomperts, S. N., Yamamoto, J. & Wilson, M. A. Neural representation of spatial topology in the rodent hippocampus. Neural Computation 26, 1–39 (2013).

32. Williams, A. H. et al. Unsupervised discovery of demixed, low-dimensional neural dynamics across multiple timescales through tensor component analysis. Neuron 98, 1099–1115.e8 (2018).

33. Sussillo, D., Jozefowicz, R., Abbott, L. & Pandarinath, C. LFADS - latent factor analysis via dynamical systems. arXiv (2016). https://arxiv.org/abs/1608.06315.

34. Wei, Z., Inagaki, H., Li, N., Svoboda, K. & Druckmann, S. An orderly single-trial organization of population dynamics in premotor cortex predicts behavioral variability. bioRxiv (2018). https://www.biorxiv.org/content/ early/2018/07/25/376830.

35. Churchland, M. M. et al. Neural population dynamics during reaching. Nature 487, 51–56 (2012).

36. Cunningham, J. P. & Yu, B. M. Dimensionality reduction for large-scale neural recordings. Nature Neuroscience 17, 1500–1509 (2014).

37. Fox, S. E. & Ranck Jr, J. B. Improved method for measuring the topological dimensions of neuronal firing rate space. Program No. 616.12. In 2017 Neuroscience Meeting Planner (Washington, DC: Society for Neuroscience, 2017). Online.

